# Drug-Target Interactions Prediction at Scale: the Komet Algorithm with the LCIdb Dataset

**DOI:** 10.1101/2024.02.22.581599

**Authors:** Gwenn Guichaoua, Philippe Pinel, Brice Hoffmann, Chloé-Agathe Azencott, Véronique Stoven

## Abstract

Drug-target interactions (DTIs) prediction algorithms are used are various stages of the drug discovery process. In this context, specific problems such as de-orphanization of a new therapeutic target, or target identification of a drug candidate arising from phenotypic screens require large-scale predictions across the protein and molecule spaces. DTI prediction heavily relies on supervised learning algorithms that use known DTIs to learn associations between molecule and protein features, allowing for the prediction of new interactions based on learned patterns. The algorithms must be broadly applicable to enable reliable predictions, even in regions of the protein or molecule spaces where data may be scarce. In this paper, we address two key challenges to fulfil these goals: building large, high-quality training datasets and designing prediction methods that can scale, in order to be trained on such large datasets. First, we introduce LCIdb, a curated, large-sized dataset of DTIs, offering extensive coverage of both the molecule and druggable protein spaces. Notably, LCIdb contains a much higher number of molecules than publicly available benchmarks, expanding coverage of the molecule space. Second, we propose Komet (Kronecker Optimized METhod), a DTI prediction pipeline designed for scalability without compromising performance. Komet leverages a three-step framework, incorporating efficient computation choices tailored for large datasets and involving the Nyström approximation. Specifically, Komet employs a Kronecker interaction module for (molecule, protein) pairs, which efficiently captures determinants in DTIs, and whose structure allows for reduced computational complexity and quasi-Newton optimization, ensuring that the model can handle large training sets, without compromising on performance. Our method is implemented in open-source software, leveraging GPU parallel computation for efficiency. We demonstrate the interest of our pipeline on various datasets, showing that Komet displays superior scalability and prediction performance compared to state-of-the-art deep learning approaches. Additionally, we illustrate the generalization properties of Komet by showing its performance on an external dataset, and on the publicly available *ℒℌ* benchmark designed for scaffold hopping problems. Komet is available open source at https://komet.readthedocs.io and all datasets, including LCIdb, can be found at https://zenodo.org/records/10731712.

## 1 Introduction

Most marketed drugs are small molecules that interact with a protein, modulating its function to prevent disease progression. Therefore, many problems in drug design boil down to the characterization of drug-target interactions (DTIs), including de-orphanizing a new therapeutic target, finding the target of a drug identified in a phenotypic screen, optimising the structure of a drug candidate to improve its ADME profile, predicting drug interaction profiles to anticipate unexpected off-targets that may lead to unwanted side effects or offer drug repositioning opportunities, or solving scaffold hopping cases.

Many different computational methods have been proposed for DTI prediction, and none can claim to help best solve all problems encountered in the drug discovery process. Indeed, they rely on very diverse principles, and on the availability of various types and amounts of data. For drug optimization problems at late stages, when many ligands have been identified or when the 3D structure of the target is available (ideally, in complex with a ligand), QSAR or structure-based virtual screening, including docking, are very efficient approaches (see reviews^1,2^). However, these approaches do not directly apply to problems such as identifying off-targets, de-orphanizing phenotypic drugs or new targets, or solving scaffold hopping cases without 3D structure. In the present paper, we tackle the design of computational models that address these categories of problems. These models need to be broadly applicable, to allow large-scale predictions in the chemical and protein spaces, even in regions of these spaces where data points about the question of interest may be scarce. Among current computational approaches, we focus on chemogenomic DTI prediction methods based on Machine Learning (ML), i.e. methods that predict whether a (molecule, protein) pair interacts or not, given known DTIs in a reference database of interactions. We formulate DTI prediction as a classification problem: (molecule, protein) pairs are classified as interacting (i.e. positive examples, labelled +1) or not interacting (i.e. negative examples, labelled -1). Indeed, chemogenomic methods offer a global framework applicable at large scales in the molecule and protein spaces to predict drugs’ protein interaction profiles or proteins’ molecular interaction profiles. The former can help identify unexpected off-targets, while the latter can help solve scaffold hopping problems.^3^

Enhancing the performance of DTI predictions at large scales requires using ever-larger training datasets and developing ML algorithms capable of scaling to these dataset sizes. Therefore, we tackle these challenges by presenting a curated large-sized dataset LCIdb and Komet, a GPU-friendly DTI prediction pipeline. These two components complement each other, resulting in state-of-the-art performance achieved with minimal use of computer resources.

## 2 State-of-the-art in chemogenomic approaches

Most chemogenomic DTI prediction methods rely on the global framework comprising three main steps presented in Figure 1. Therefore, we shortly review state-of-the-art approaches used in these three steps.

**Figure 1:**
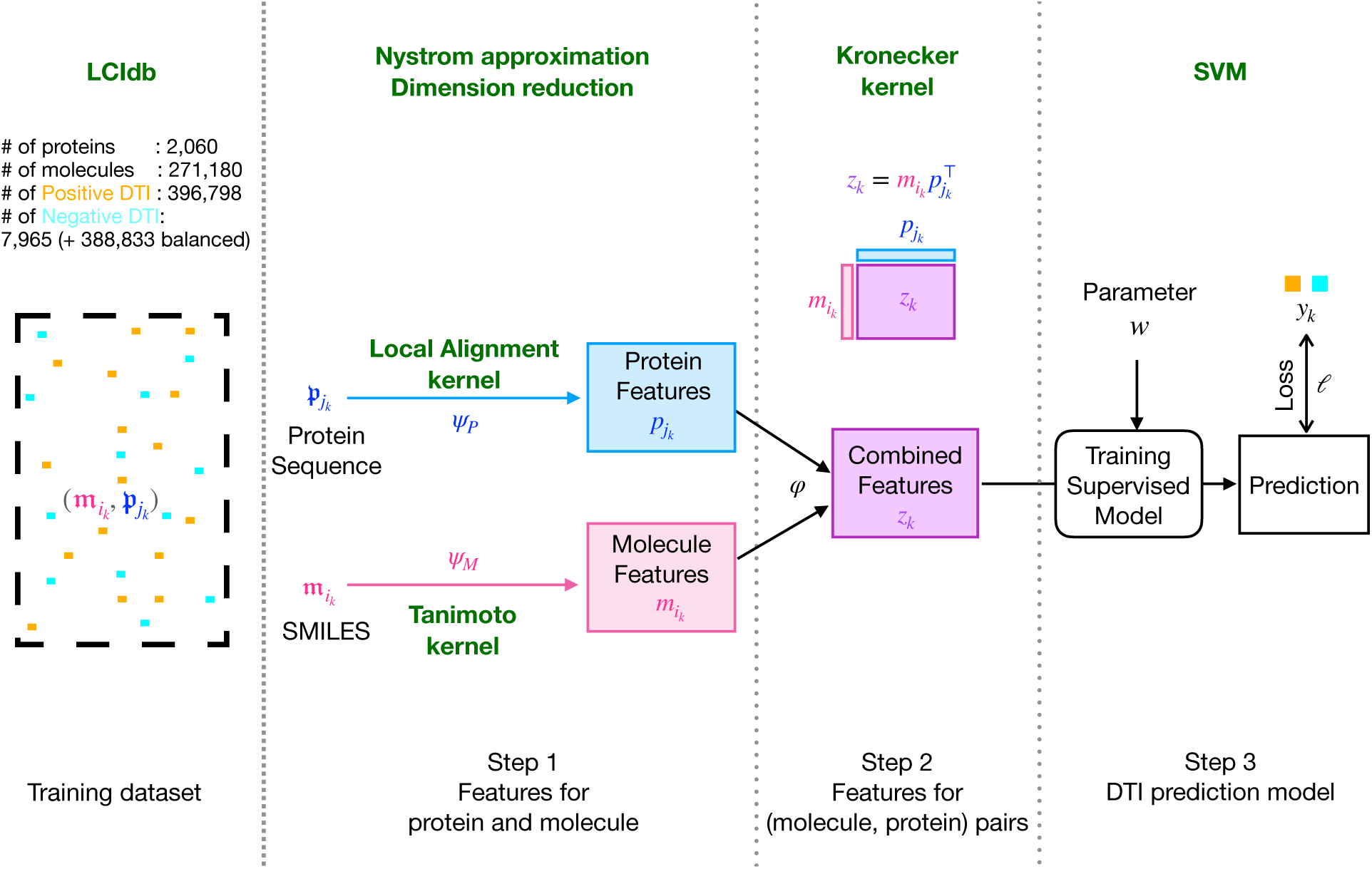
Komet’s global framework for DTI prediction. Specific aspects of Komet itself are in bold green, while generic elements of such as the typical 3-step framework are in black.

### 2.1 Step 1: Molecule and protein Features

Various methods have been designed to compute feature representations for proteins and molecules.^4^ We will focus here on features that are applicable at large scales, and therefore, that rely on the 2D chemical structures of molecules and primary sequences of proteins.

For molecules, several types of 2D structure encodings are considered, as discussed in recent papers.^5,6^ They can globally be classified into: (1) string-based formats such as the Simplified Molecular-Input Line-Entry System (SMILES),^7^ or the International Chemical Identifier (InChI);^8^ (2) table-based formats that represent the chemical graph of the molecule, such as the SDF format.^9^ From these primary encodings, various molecular features (also called descriptors) can be derived, including: (1) feature-based vectors encoding various molecular characteristics, such as Morgan fingerprints or Extended-Connectivity Fingerprints (ECFP),^10^ as well as 2D and 3D pharmacophore fingerprints as described in the RDKit toolbox^11^; (2) computer-learned features derived by neural networks in deep learning approaches. These features can be learned for example from recurrent neural networks or convolutional neural networks that use SMILES strings as input.^12,13^ Graph convolutional networks have also been applied to 2D molecular graphs to learn small molecule features,^14,15^ and strategies to pre-train graph neural networks have been studied by^16^ to compute molecule features. Similar to natural language models, Mol2vec^17^ and SMILES2vec^18^ adapt the principles of the word2vec method^19^ to learn features for molecular structures. Additionally, transformer-based models like MolTrans^20^ have emerged in this domain. Finally, other learned representation methods such as X-Mol^21^ or MolGNet^22^ use AutoEncoder (AE) techniques to compute molecular features.

Similarly, the primary structure of proteins can globally be described by: (1) string-based representations corresponding to their primary sequence of amino acids; (2) vector-based feature representations, where the elements of the vector are features that are calculated according to various protein characteristics, as reviewed in Zhu et al. ^23^. They include the classically-used composition, transition, and distribution (CTD) descriptors;^24^ (3) computer-learned descriptors derived by neural networks in deep learning approaches. In this context, protein features can be acquired by a variety of deep learning architectures, including recurrent neural networks or convolutional neural networks,^12,13^ as well as transformer models.^20^ As in natural language models, protein features can also be learned from pre-trained transformer-based models on external tasks such as ESM2,^25^ or auto-encoder models such as ProtBert^26^ and ProtT5XLUniref50.^26^

### 2.2 Step 2: Features for (molecule, protein) pairs

The second step of many DTI prediction pipelines consists of defining a representation for (molecule, protein) pairs, thus defining a latent space for pairs. The method that is used to define this latent space has a critical impact on the prediction performance, and a key aspect is that the features representing the (molecule, protein) pair should capture information about the interaction, which is not fully achieved by simple concatenation between molecule and protein features.^27^ Therefore, step 2 usually consists of a non-linear mixing of the protein and molecule features, to better encode information about interaction determinants. One common approach is to use the tensor product, which is equivalent to a Kronecker kernel.^28,29^ Alternatively, in deep learning methods, the features for pairs can be learned from an interaction module that consists of fully connected multi-layer perceptrons.^12,30–32^ Attention mechanisms applied to molecule and protein features constitute another option.^13,20,33^ Then, the last layer of the network can be interpreted as a feature vector representing the (molecule, protein) pairs.

### 2.3 Step 3: DTI prediction model

The third step consists of a supervised classifier that is trained in the latent space of (molecule, protein) pairs, using a training dataset of positive and negative DTIs. These classifiers include tree-based methods^34^ and network-based inference approaches.^35^ In linear models, step 3 consists of the optimization of the weights applied to the pair features calculated in step 2, according to a logistic loss, or a hinge loss for Support Vector Machines (SVM).^36^ For example, all methods of Pahikkala et al. ^29^, Nagamine and Sakakibara ^37^, Jacob and Vert ^38^, Playe et al. ^39^ rely on a linear model in a latent representation of pairs. In deep learning chemogenomic algorithms, step 3 relies on the pair features determined by the last layer of the neural network in step 2. The features’ weights are optimized based on a loss function, typically binary cross-entropy, as the input progresses through the network in a feed-forward manner. This approach is used in several recent papers.^12,13,20,30–33^

### 2.4 Challenges in chemogenomic studies

Although different chemogenomic approaches have been proposed, as briefly reviewed above, all require a training dataset of positive and negative (molecule, protein) pairs. Recent ML chemogenomic algorithms have often been trained on small to medium-sized benchmarks that present various biases. Indeed, most classical benchmark datasets are extracted from a single biological database, and often favour drug and target families that have been more widely studied, and for which many known DTIs have been recorded.^40,41^ Additionally, Bagherian et al. ^42^ highlights that most datasets use negative DTIs randomly chosen among pairs with unknown interaction status, and may therefore include false negative DTIs. One suggestion to overcome this problem is to derive training datasets from interaction databases that compile continuous values for binding affinities, and choose stringent activity thresholds to derive confident positive and negative pairs, as suggested by Wang et al. ^43^.

In addition, learning chemogenomic models that are broadly applicable and can generalize to many different families of proteins and drugs require training on very large, high-quality, verified and well-established DTI datasets. This appears to be an important bottleneck since publicly available training datasets that meet these criteria are seldom.

However, training ML algorithms on very large datasets, potentially comprising hundreds of thousands of molecules, and therefore of DTIs, leads to challenges in terms of computation times and memory requirements. In particular, the choice of the interaction module in step 2 has significant implications for computation time and memory resources in large-sized datasets. In the case of deep learning approaches, the complexity of neural network architectures, and the size of parameter spaces, may also contribute to the computational expense. Learning the interaction module requires iteratively adjusting the model parameters, leading to time-consuming training phases.

Overall, there is a critical need for chemogenomic approaches that are computationaly frugal so that they can scale to very large datasets.

## 3 Contributions

### 3.1 Global organisation of the paper

In the present paper, we tackle the two important challenges mentioned above, which are critical when the goal is to make large-scale predictions in the protein and molecule spaces:

- in Section 4.2, we build the Large Consensus Interaction dataset, called LCIdb hereafter, a new, very large, high-quality dataset of DTIs that was designed to train chemogenomic ML algorithms for large-scale DTI prediction. In particular, LCIdb comprises a much larger number of molecules than commonly used datasets, offering a better coverage of the chemical space. Additionally, we paid attention to limiting potential bias among negative DTIs.
- in Sections 4.3 and 4.4, we propose Komet (Kronecker Optimized METhod), an efficient DTI prediction method that lies within the global pipeline presented in Figure 1. This method incorporates specific encoding and computation choices that provide scalability for very large training datasets, without compromising prediction performance.

We show that Komet competes with or outperforms state-of-the-art deep learning approaches for DTI prediction on medium-sized datasets, but scales much better to very large datasets in terms of prediction performances, computation time, and memory requirements (see Section 5.4).

Finally, we illustrate the performance of Komet trained on LCIdb using DrugBank as an external dataset for DTI prediction, and on a publicly available benchmark^44^ designed to evaluate the performance of prediction algorithms in solving difficult scaffold hopping problems.

Komet adopts the global three-step framework shown in Figure 1, which aligns with recent computational pipelines, such as in Huang et al. ^30^. However, Komet includes specific choices shown below, while mathematical details are provided in Materials and Methods.

### 3.2 Principle of the Komet pipeline

Komet derives from kernel SVMs because these algorithms display good generalisation properties and are computationally efficient,^39^ which are two important characteristics for large-scale DTI prediction. In this framework, a classical method to build a kernel for (molecule, protein) pairs is to use the Kronecker product of a molecule kernel and a protein kernel. This approach is well suited to small datasets, but becomes computationally untractable when the size of the DTI training set becomes very large, because the resulting kernel matrix cannot be stored in memory. To get around this limitation, we chose to go back to working with features by computing explicit feature maps for the molecules, proteins, and pairs kernels. In other words, our features are such that their dot product is equal to the chosen kernel. We can then train linear SVMs, which for large training sets can be done much more efficiently than kernel SVMs, in this new feature space.

In step 1, molecule features are computed by factorization of the molecule kernel *k_M_*, so as to match the kernel’s feature map: for any two molecules **m** and **m**′, their feature vectors *ψ_M_* (**m**) and *ψ_M_* (**m**′) are such that *⟨ψ_M_* (**m**)*, ψ_M_* (**m**′)⟩ = *k_M_* (**m**, **m**′). Protein features are similarly computed from the protein kernel. Due to the large number of molecules in LCIdb, this decomposition is computationally expensive for the molecule kernel. Therefore, we use the Nyström approximation^45,46^ to compute an approximation of the molecule kernel’s feature map that only uses a small, randomly chosen set of *m_M_* landmark molecules from the training set to obtain molecule feature vectors of size *m_M_*. In addition, computations involved in the Nyström approximation include the diagonalisation of the small kernel matrix restricted to the landmark molecules (a matrix of size *m_M_ × m_M_*) using single value decomposition. This offers the possibility to further reduce the dimension of molecule feature vectors to a size of *d_M_ ≤ m_M_*. The impact on Komet’s prediction performances and computation requirements of the two parameters, i.e. the numbers *m_M_* of molecule landmarks and the dimension *d_M_* of the molecule feature vectors, is studied in Section 5.2. Because the number of proteins in LCIdb is rather small, we do not apply these approximations to the computation of protein features.

In step 2, the interaction module consists of the tensor product between the protein and molecule feature vectors. The size of the resulting feature vector representing (molecule, protein) pairs, which is *d_M_ d_P_* (where *d_P_* is the number of proteins in the training set), can be prohibitively large in terms of computation time and memory. However, solving the SVM in step 3 only involves the dot products between (molecule, protein) pairs. Thanks to classical factorization properties of tensor products, this allows to solve the SVM while avoiding the explicit calculation of the pairs’ feature vectors, thus addressing the challenges posed by large datasets. Another motivation for using the tensor product is that it offers a systematic way to encode (molecule, protein) pairs, independently of the choice of molecule and protein features. Furthermore, the tensor product of vectors, whose descriptors are all possible products between a protein descriptor and a molecule descriptor, is exhaustive and may capture key determinants that govern (molecule, protein) interactions.^47^ As observed in Section 5, this tensor product representation efficiently captures information about (molecule, protein) binding.

In step 3, Komet uses a simple SVM loss together with, a full batch, BFGS optimization algorithm. This allows to leverage the Kronecker factorization of pairs’ features, leading to a significant speedup of the training. It is important to note that, in the proposed approach, steps 2 and 3 are executed simultaneously. This is made possible by avoiding the implicit calculation of pairs’ features, thanks to the Kronecker interaction module.

Our method is implemented in an open source software, leveraging parallel computation on GPU through a PyTorch^48^ interface, and is available at https://komet.readthedocs.io. All datasets, including LCIdb, can be found at https://zenodo.org/records/10731712.

## 4 Materials and Methods

We first recall known and publicly available medium-sized DTI datasets that are used in the present paper (Section 4.1), and describe the construction of our large-sized DTI dataset LCIdb (Section 4.2). Then, we detail our computational approach for large-sized DTI prediction with Komet (Sections 4.3 and 4.4), and present the methodology used to compare the performance of Komet to those of a few state-of-the-art deep learning algorithms (Section 4.5). Finally, we introduce *ℒℌ*, a publicly available benchmark dataset for scaffold hopping problems.

### 4.1 Medium-scale datasets

We first use medium-scale datasets to compare the performance of Komet to those of state-of-the-art algorithms: BIOSNAP, BIOSNAP_Unseen_drugs, BIOSNAP_Unseen_proteins, BindingDB, and DrugBank. The four first of these datasets are publicly available and were established in Huang et al. ^20^. They are used in various recent studies.^30,49^ The last one is the DrugBank-derived dataset established in Najm et al. ^50^, from which we built an additional set called DrugBank (Ext) to be used as an external validation dataset, as detailed below.

We perform a 5-fold cross-validation with complete replacement of the test dataset. In Supplementary Table S2, we also evaluate performance on a train/validation/test split of the data, as done in Huang et al. ^20^ and Singh et al. ^49^.

#### BIOSNAP in its three prediction scenarios

The ChGMiner dataset from BIOSNAP^51^ contains exclusively positive DTIs. Negative DTIs are generated by randomly selecting an equal number of positive DTIs, assuming that a randomly chosen (molecule, protein) pair is unlikely to interact. As proposed in Huang et al. ^20^, we considered three scenarios that are achieved based on different splits of BIOSNAP to build the different folds.The first scenario, referred to as BIOSNAP, corresponds to a random splitting of the DTIs in BIOSNAP. In the BIOSNAP_Unseen_targets scenario, the Train and Test sets do not share any protein. The BIOSNAP_Unseen_drugs dataset follows a similar process for molecules. The two last scenarios allow us to evaluate the generalization properties of the algorithm on proteins or molecules that were not seen during training.

#### BindingDB-derived dataset

The BindingDB database^52^ stores (molecule, protein) pairs with measured bioactivity data. We used a dataset derived from BindingDB and introduced by Huang et al. ^20^, where BindingDB is filtered to include only pairs with known dissociation constants (Kd). Pairs with Kd < 30 nM are considered positive DTIs, while those with Kd > 30 nM values are considered negative. This leads to a much larger number of negative DTIs than positive DTIs. Although the resulting dataset does not include the whole BindingDB database, for the sake of simplicity, it will be called BindingDB hereafter.

#### DrugBank-derived datasets

We used the dataset provided in Najm et al. ^50^. This dataset was built by filtering drug-like molecules and human protein targets in the DrugBank database,^53^ adding an equal number of negative DTIs through balanced sampling. More precisely, to avoid bias towards well-studied proteins for which many interactions are known, negative examples are randomly chosen among unlabeled DTIs in such a way as to ensure that each protein and each drug appear an equal number of times in positive and negative interactions, using a greedy algorithm. This dataset will be referred to as DrugBank in the following, for the sake of simplicity, and corresponds to the dataset called DrugBank (S1) in the original paper.

We created another dataset called DrugBank (Ext), derived from the above dataset, and used it as an external validation to compare the prediction performances of the considered algorithms when trained on BindingBD or on LCIdb. Positive interactions from DrugBank were selected, excluding those present in BindingDB and LCIdb, to gather a set of positive DTIs absent from the BindingDB and LCIdb datasets. All other DTIs in DrugBank are kept in DrugBank (Ext). As above, balanced negative interactions were added in DrugBank (Ext), using the greedy algorithm of Najm et al. ^50^.

### 4.2 Building the new large-scale dataset LCIdb

To build a large-sized dataset of DTIs, we started from the Consensus database described by Isigkeit et al. ^54^, as it combines and curates data from prominent databases including ChEMBL,^55^ PubChem,^56^ IUPHAR/BPS,^57^ BindingDB,^58^ and Probes & Drugs.^59^ Compounds in the Consensus dataset are already standardized as described in Isigkeit et al. ^54^, and therefore, we relied on this standardization. It involves the removal of salts, the creation of canonical SMILES, the canonicalization of tautomers, and molecular structure checks, ensuring consistent structural information across all entries. We then filtered the DTIs in this database according to four criteria, as detailed below.

#### Filtering positive DTIs

1. Chemical structure quality filter: for DTIs present in several of the source databases, we only retained those for which the SMILES representation of the molecule was identical in all sources, to exclude potential erroneous (molecule, protein) pairs. We only kept molecules with molecular weights between 100 and 900 g.mol^-1^, which is a standard choice for selecting drug-like molecules. Among these molecules, we selected those that target at least one human protein. These filters were used because the goal was to build a training dataset of DTIs that are relevant in the context of drug discovery projects.
2. Bioactivity filter: we retained only DTIs for which the negative logarithm of inhibition constant Ki, dissociation constant Kd, or half maximal inhibitory concentration IC50 measurements were available in at least one of the source databases.
3. Quantitative bioactivities filter: for DTIs with bioactivity measurements present in multiple source databases, we only retained those with bioactivities within one log unit from one another.
4. Binary labelling of DTIs: Bioactivity measurements (first Kd, then Ki, then IC50) were converted into binary interactions based on a threshold. When multiple bioactivity measurements have a difference of less than one log unit: if the average bioactivity value was less than 100 nM (10^-7^M), the interaction was labelled as a positive DTI (binding). If the average bioactivity value was greater than 100*µ*M (10^-4^M), the interaction was labelled as a negative DTI (non-binding). If the average bioactivity value was in the intermediate range, i.e. between 100 nM and 100*µ*M, DTIs were labelled as non-conclusive. When multiple bioactivity measurements differ by more than one logarithmic unit: if all bioactivity values were less than 100 nM (10^-7^ M), the interaction was classified as positive (binding). If all bioactivity values were greater than 100 *µ*M (10^-4^ M), the interaction was classified as negative (non-binding). If the bioactivity values were between 100 nM and 100 *µ*M, the interaction was classified as known non-conclusive.

This scheme leads to the selection of 271 180 molecules, 2 060 proteins, 396 798 positive interactions and 7 965 negative interactions. We then added negative interactions to build a balanced dataset, as described below. Figure 2 illustrates the filters applied on the considered databases to build LCIdb.

**Figure 2:**
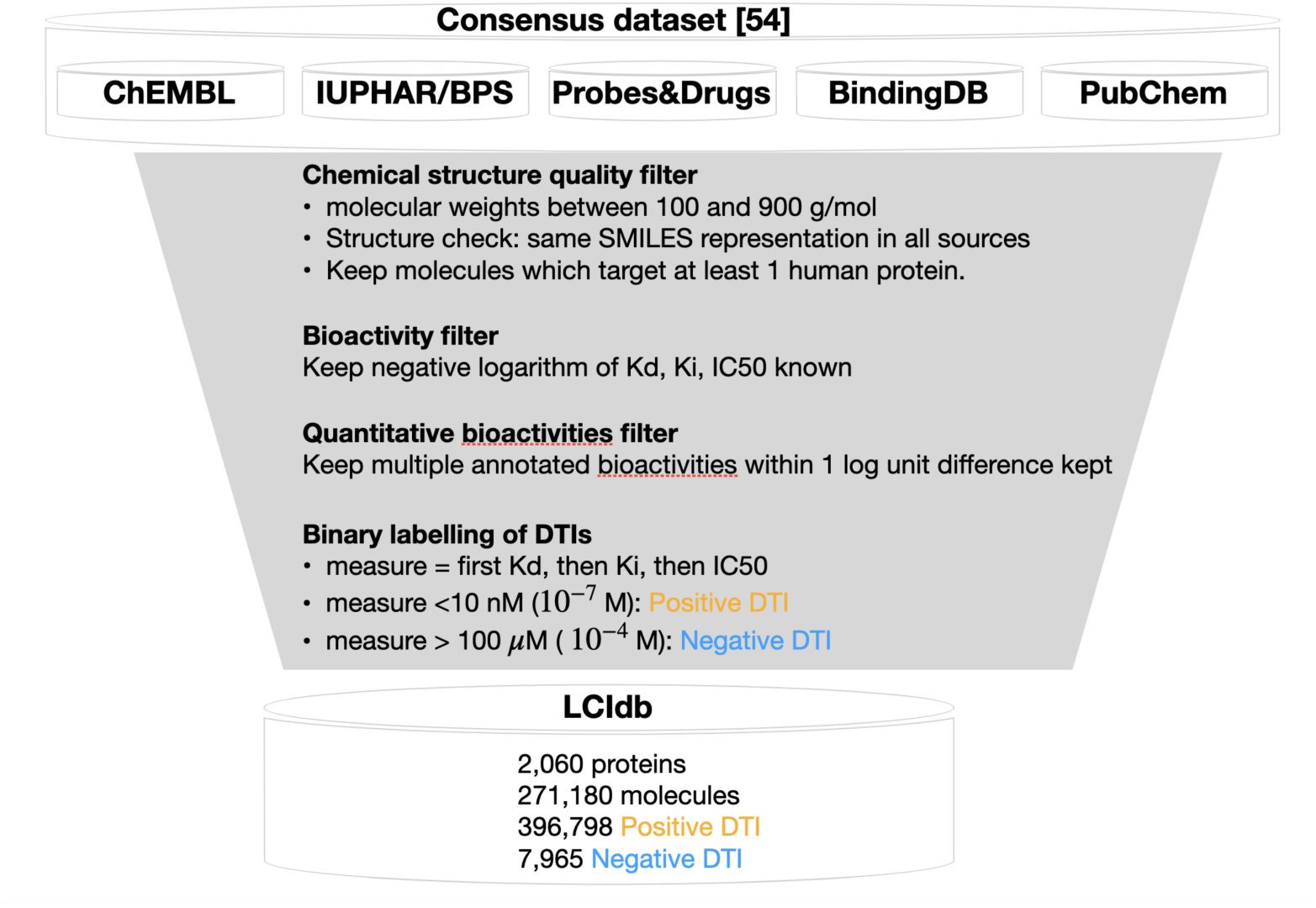
Flowchart describing the successive filters applied on the considered databases to build LCIdb.

#### Completion of a balanced negative DTI dataset

We perform a 5-fold cross-validation with complete replacement of the test dataset. We used unlabeled DTIs to include negative interactions in these sets, assuming most unknown DTIs are negative. For the training set, the selection of additional negative interactions should be designed with care to tackle two classical issues: (1) reduce the number of false negative DTIs present in the training set; (2) correct potential statistical bias in the database towards highly studied molecules or proteins. To take into account the former, we excluded known non-conclusive interactions, and for the latter, we applied the algorithm by Najm et al. ^50^ for selecting additional negative DTIs. In the test sets, remaining negative and randomly chosen unknown interactions are added. These sets form LCIdb, mirroring the DrugBank dataset scenario discussed in Section 4.1.

#### Different prediction scenarios

To evaluate performance in different prediction scenarios, we also derive different datasets from to LCIdb based on specific splits of the Train and Test sets, as proposed in Huang et al. ^20^ and Singh et al. ^49^. Datasets are built to correspond to LCIdb, LCIdb_Unseen_drug, LCIdb_Unseen_protein, and LCIdb_Orphan (unseen molecule and protein) scenarios. We added the Orphan case, which presents the greater difficulty for prediction tasks.

More precisely: (1) LCIdb is balanced in positive and negative pairs chosen at random; (2) LCIdb_Unseen_drugs is built so that (molecule, protein) pairs in one of the Train/Test sets only contain molecules that are absent from the other set; (3) LCIdb_Unseen_targets is built so that (molecule, protein) pairs in one of the Train/Test sets only contain proteins that are absent from the other set; (4) LCIdb_Orphan is built so that (molecule, protein) pairs in one of the Train/Test sets only contain proteins and molecules that are absent from the other set. The number of drugs, targets, and interactions in these four datasets is given in Table 1.

**Table 1:** Numbers of molecules, proteins, and positive/negative DTIs in the considered datasets. “random” indicates that negative DTIs were randomly chosen among unlabeled DTIs. “balanced” indicates that negative DTIs were randomly chosen among unlabeled DTIS, but in such a way that each protein and each drug appears in the same number of positive and negative DTIs.

| Datasets | Molecules | Proteins | Positive DTIs | Negative DTIs |
| --- | --- | --- | --- | --- |
| BIOSNAP | 4,510 | 2,181 | 13,836 | (13,647 random) |
| Unseen_drugs |  |  | 13,836 | (13,647 random) |
| Unseen_targets |  |  | 13,836 | (13,647 random) |
| BindingDB | 7,161 | 1,254 | 9,166 | 23,435 |
| DrugBank | 4,813 | 2,507 | 13,715 | (13,715 balanced) |
| DrugBank (Ext) | 4,257 | 1,216 | 10,838 | (10,838 balanced) |
| LCIdb | 271,180 | 2,060 | 396,798 | 7,965 (+ 388,833 balanced) |
| Unseen_drugs | 271,180 | 2,060 | 396,798 | 7,965 (+ 388,833 balanced) |
| Unseen_targets | 271,180 | 2,060 | 396,798 | 7,965 (+ 388,833 balanced) |
| Orphan | 191,901 | 2,060 | 208,041 | 7,965 (+ 200,076 balanced) |

### 4.3 Features for proteins and molecules in Komet

As introduced in Section 3.2, the initial step of our DTI prediction framework consists of computing simple and fixed features for molecules and proteins, based on kernels for proteins and molecules. Therefore, these kernels are presented below, and then we provide mathematical details about feature calculations.

#### Choice of molecule and protein kernels

The feature maps *ψ_M_* and *ψ_P_* depend on the choice of molecule and protein kernels. We follow the choices made in Playe et al. ^39^ and adopt the Tanimoto kernel *k_M_* for molecules. For each molecule **m** represented in SMILES format, we calculate ECFP4 fingerprints, generating a 1024-bit binary vector using the RDKit^60^ package. Values of the Tanimoto kernel between two molecules are then computed as the Jaccard index between their fingerprints. The Tanimoto kernel hence measures the similarity between two molecules based on the substructures they share. Based on proteins represented by their primary sequence **p** of amino acids, we opt for the Local Alignment kernel (LAkernel).^61^ In the context of DTI prediction at large scales, requiring to measure similarity between proteins belonging to distant families in terms of evolution, this kernel *k_P_* appeared to be relevant because it detects remote homology by aggregating contributions from all potential local alignments with gaps in the sequences, thereby extending the Smith-Waterman score.^62^ For both kernels, we used the same hyperparameters as Playe et al. ^39^, where they were adjusted by cross-validation.

#### Computing molecule and protein features from molecule and protein kernels

In Komet, the feature vector *ψ_M_* (**m**) for a molecule **m** is computed using an explicit feature map for the molecule kernel *k_M_*. *k_M_* (**m**, **m**′) can be viewed as a similarity measure between two molecules. The explicit feature map for *k_M_* is computed by factorization of the empirical kernel matrix *K_M_* ∈ ℝ*^n_M_ × n_M_^*, where (*K_M_*)*_i,j_* := *k_M_* (**m***_i_*, **m***_j_*) for all molecules in the training set. This factorization can be written as 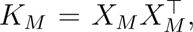 where *X_M_* is a matrix where the *i*-th line is the explicit feature map of **m***_i_*, the *i*-th molecule of the training set. However, given the number of molecules in LCIdb (*n_M_* = 271, 180), and therefore the size of *K_M_*, the factorization of this kernel matrix leads to computation and memory burdens. Therefore, we instead leverage the Nyström approximation^45,46^ to efficiently compute molecule features using an approximate feature map. More precisely, we use a small set of landmark molecules 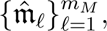 with *m_M_ ≪ n_M_*, that are randomly chosen in the training dataset. Then, we compute a small kernel matrix over these landmark molecules: 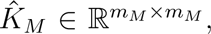 where 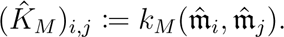 We define the extrapolation matrix *E* ∈ ℝ*^m_M_ × m_M_^* from the Singular Value Decomposition (SVD) of 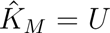 diag(*σ*)*U*^T^ as *E* := *U* diag(*σ*)^-1/2^. This extrapolation matrix allows to compute molecule feature vectors for any molecule **m** (in particular for molecules in the training set that are not in the landmark set) as:

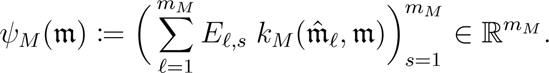

Note that these feature vectors satisfy the relation 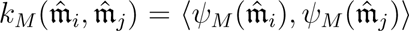 for the landmark molecules (see Section 5 of Supporting Information for details).

For any molecule **m** that is not in the landmark set, 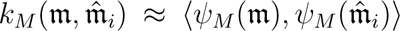 (see Section 5 of Supporting Information for details). Hence, *E* allows us to “extrapolate” the feature map *ψ_M_*, which is an explicit feature map of *k_M_*, from the landmarks to new molecules.

Furthermore, we can reduce the dimension of the feature vectors by keeping in *E* only the *d_M_* (*d_M_ ≤ m_M_*) higher eigenvalues in the SVD of 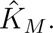 We then define

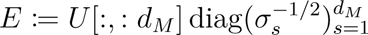

and have

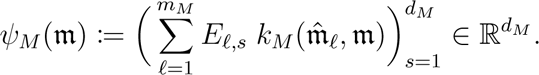

Finally, we mean-center and normalize the features:

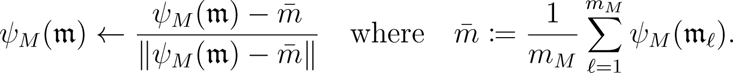

Note that the “best way” to choose landmark molecules corresponds to a random choice, to sample the chemical space as uniformly as possible. Randomly picking “enough” landmark molecules is a way to sample the chemical diversity present in the training set, without introducing bias, allowing a better extrapolation of the non-landmark molecules’ features.

We adopt a similar approach to build feature vectors for proteins. However, the number of proteins in LCIdb being much smaller than that of molecules (*n_P_* = 2 060), an explicit feature map for the protein kernel can be computed exactly, using kernel factorization (by SVD), without resorting to Nyström approximation or dimension reduction. Therefore, in the case of proteins, *d_P_* = *m_P_* = *n_P_*.

### 4.4 Large-scale chemogenomic framework with Komet

We address DTI prediction as a supervised binary classification problem, incorporating established steps, as outlined in Sections 2.2 and 2.3.

#### Features for molecule-protein pairs

Let us consider a DTI dataset containing molecules and proteins 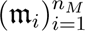 and 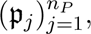 where *n_M_* and *n_P_* are respectively the number of molecules and proteins in the dataset. To alleviate notations, in what follows, we denote by *m* := *ψ_M_* (**m**) the feature vector of a molecule **m** and by *p* := *ψ_P_* (**p**) the feature vector of a protein **p**.

The training dataset consists of a set of *n_Z_* (molecule, protein) pairs with indices 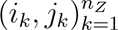 and their associated labels *y_k_* ∈ {-1, 1}. If *y_k_* = 1 (resp. -1), molecule **m***_i_k__* and protein **p***_j_k__* interact (resp. do not interact). The classification is performed in the space of pairs, which we define as the tensor product of the space of molecules and the space of proteins.

Hence, the feature vector for a pair **m, p** is given by *φ*(**m, p**) := (*m*[*s*]*p*[*t*])_1≤*s* ≤ *d_M_*, 1≤*t* ≤ *d_P_*_ *∈* ℝ*^d_Z_^*, where *m*[*s*] is the *s*-th coordinate of *m* and *p*[*t*] is the *t*-th coordinate of *p*.

Thus, the space of pairs has dimension *d_Z_* = *d_M_ d_P_*. These features correspond to the use of a Kronecker kernel, already shown to be efficient in kernel-based chemogenomic approaches.^29,39,50^ Using a Kronecker kernel is crucial in our approach, not only because it is a state-of-the-art method, but also due to its favourable mathematical properties, which we will detail below. It is worth noting that our approach avoids explicitly calculating the feature map *φ*, which mitigates the computational burden associated with the large value of *d_Z_*.

#### SVM classification

Our classification approach follows previous work (see Section 2.3), relying on a linear model with weight vector *w* ∈ ℝ*^d_Z_^* and bias term *b* ∈ ℝ. The class decision for a pair feature vector *z* ∈ ℝ*^d_Z_^* is determined by sign(〈*w, z*〉 + *b*) *∈* {-1, 1}. The parameters *w* and *b* are obtained by minimizing a penalized empirical risk:

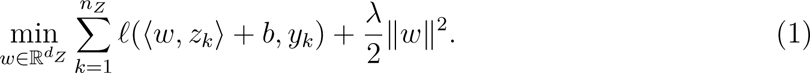

In Komet, we employ a Support Vector Machine (SVM) classification where *ℓ*(*y′, y*) = max(0, 1 *- yy′*).

The minimization of Equation (1) is computationally demanding, particularly when *n_Z_* and *d_Z_* are large. A conventional Stochastic Gradient Descent (SGD)^63^ can result in slow convergence. Therefore, we use an alternative approach that leverages the specific structure of our feature map *φ*, as was previously done by Airola and Pahikkala ^64^. Specifically, we exploit: (1) the tensor product nature of *φ* and (2) the fact that the sizes *n_M_* and *n_P_* of the input databases are much smaller than the number *n_Z_* of interactions.

#### Efficient computation

The core ingredient leading to a significant improvement in computational efficiency on a large-sized dataset is the efficient computation of the gradient by bypassing the evaluation of *φ*. Indeed, the function to be minimized in Equation (1) has the form 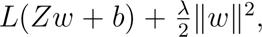 where the rows of *Z* ∈ ℝ*^n_Z_ ×d_Z_^* are the vectors 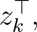, and *L* takes into account *ℓ* and *y*. The main computational burden for evaluating this function and its gradient is the computation of *Zw*. A naive implementation would require *n_Z_d_Z_* operations just to compute *Z*, which would be unavoidable if one used a generic *φ*, such as a deep neural network. However, we bypass this bottleneck by directly computing *Zw*. This relies on the following identity:

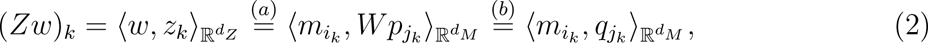

where *W* ∈ ℝ*^d_M_ × d_P_^* is such that it has *w* as flattened representation in ℝ*^d_Z_^* and *q_j_*:= *W_p_j__*.

Equality (a) exploits the tensor product structure of *φ*. Please refer to Section 6 of Supporting Information for details for detailed proof.

Equality (b) is interesting because all the 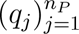 can be computed in only *n_P_ d_Z_* operations. Once this has been computed, evaluating all *n_Z_* values of (*Zw*)*_k_* = *〈m_i_, q_j_〉*_R_*d_M_* require *n_Z_d_M_* operations. We then minimize Equation (1) using a full batch method, which enables the use of efficient quasi-Newton methods. In practice, we use the BFGS method with limited memory.^65^ The complexity of our algorithm is then *O*(*n_P_ d_Z_* + *n_Z_d_M_*) where *O*(.) takes into account the number of iterations of the BFGS algorithm to reach a fixed accuracy. This number is quite small (10 to 50) in our numerical experiments. Note that we can exchange the role of the protein features and the molecule features in this calculation, resulting in a complexity of *O*(*n_M_ d_Z_* + *n_Z_d_P_*). In our setting *n_P_ ≪ n_M_* so we prefer the initial formulation of Equation (2).

#### From classification to probability estimation

Once the weight vector *w* has been computed, Platt scaling^66^ computes a probability of belonging to the positive class using the formula *p_k_* := *σ*(*-y_k_*(*s〈z_k_, w〉* + *t*)), where *σ* is the logistic function 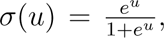 and the scale *s* (which can be interpreted as a level of confidence) and the offset *t* need to be optimized. This is achieved by minimizing the same energy as in logistic regression:

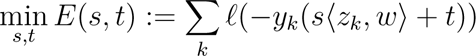

where *ℓ*(*u*) := log(1 + *e^u^*). We use the BFGS method to solve this equation.

### 4.5 Evaluation of prediction performance

Comparing the prediction performances of various algorithms requires defining the evaluation strategies and the metrics used. We formulate the DTI prediction problem as a classification task, therefore, we use AUPR (area under the precision–recall curve) as a metric to compare prediction performances of various algorithms trained on various medium or large-scale datasets. There is only one hyperparameter in our model, as shown in Equation (1). We select the best *λ* ∈ {10^-11^, 10^-10^*, …,* 10, 100} based on AUPR performance from a 5-fold cross-validation, each time with new landmark molecules and approximated molecule features.

### 4.6 Large-scale predictions in the chemical space for solving large-step scaffold hopping problems

To assess the interest of various algorithms to solve scaffold hopping problems, we used the *ℒℌ* benchmark^44^ (https://github.com/iktos/scaffold-hopping). This high-quality database comprises 144 pairs of ligands for 69 diverse proteins. These pairs constitute well-characterized and non-redundant examples of large-step scaffold hopping cases. They were extracted from the PDBbind database,^67^ which gathers crystallographic structures of proteins in complexes with various ligands, and they are composed of two highly dissimilar molecules sharing similar binding modes within the same protein pocket. As detailed in the original paper, these pairs were filtered from PDBbind according to various criteria, including Morgan and Murko-based Morgan similarities lower than 0.3 and 0.6, respectively. Globally, the subsequent filters ensured that the *ℒℌ* benchmark contains only ‘true’ large-step scaffold hopping cases, i.e. pairs of molecules that bind similarly to the same protein while displaying highly dissimilar 2D structures. In addition, for each pair, 499 decoys were carefully picked to avoid bias towards either of the two ligands. In particular, they present similar global physical and chemical properties to the ligands, while they are as distant from each ligand of the pair as these ligands are from each other, in terms of 2D chemical structure.

With the *ℒℌ* benchmark, the performances of computational methods are evaluated as follows: for each of the 144 pairs, one ligand is designated as the known active while the other is considered as the unknown active and added to the 499 decoys. Given the known active, each method ranks the unknown active among the 499 decoys, the lower the rank, the better. For each pair, one molecule or the other can be assigned as the known active, which leads to 288 scaffold hopping cases to solve.

For each case, the considered algorithms were trained with one molecule of the pair assigned as the only known active for the query protein. If the known interaction was absent from the training dataset that was used, it was added to it, and all other interactions involving the query protein potentially present in the database were removed. After training, the algorithms ranked the unknown active and the 499 decoy molecules, according to the predicted binding probabilities of the (molecule, query protein) pairs.

As in Grisoni et al. ^68^, we employ three criteria to compare ranking algorithms: (1) Cumulative Histogram Curves (CHC) are drawn to represent the number of cases where a method ranks the unknown active below a given rank, with better-performing methods having curves above others; (2) Area Under the Curve (AUC) of CHC curves provide a global quantitative assessment of the methods; (3) the proportion of cases where the unknown active was retrieved in the top 1% and 5% best-ranked molecules.

## 5 Results

In the following, we first present the new LCIdb DTI dataset, analyze its coverage of the molecule and protein spaces, and compare it to other available and widely used datasets. Next, we explore different parameters within the Komet pipeline, to find a balance between speed and prediction performance. We then show that Komet displays state-of-the-art DTI prediction performance capabilities on the considered medium- and large-sized datasets, and on the external dataset DrugBank (Ext). Finally, we highlight the efficiency of our approach on the publicly available (*ℒℌ*) benchmark dataset designed to address challenging scaffold hopping problems.

### 5.1 Coverage of the protein and molecule spaces in the LCIdb dataset

Different reviews introduce numerous biological databases that can be used to derive large-sized training datasets,^4,42^ to best cover the protein and molecule spaces. Following Isigkeit et al. ^54^, we combine and filter curated data from prominent databases including ChEMBL^55^ PubChem,^56^ IUPHAR/BPS,^57^ BindingDB,^58^ and Probes & Drugs,^59^ and built LCIdb, a large-sized high-quality DTI database, as detailed in Section 4.2. Table 1 provides the numbers of molecules, proteins, and interactions in all the DTI training datasets considered in the present study.

Table 1 reveals that DrugBank- or BIOSNAP-derived datasets and BindingDB share a few characteristics: their numbers of proteins are similar (in the range of one to two thousand), their numbers of molecules are modest (in the range of a few thousand), their number of known positive DTIs are similar (in the range of thousands). BindingDB contains true negative DTIs, while the DrugBank- or BIOSNAP-derived datasets use DTIs of unknown status as negative DTIs, randomly chosen for BIOSNAP-derived datasets, and randomly chosen in such a way that all molecules and proteins appear in the same number of positive and negative DTIs (labelled “balanced” in Table 1) for the DrugBank-derived datasets. Overall, these observations underline the need for a larger dataset, as required for chemoge-nomic studies. As shown in Table 1, LCIdb includes 40 times more molecules and 30 times more positive DTIs than the other considered datasets, the number of human proteins being in the same order of magnitude.

However, it is important to evaluate whether this larger number of molecules corresponds to better coverage of the chemical space and whether the different datasets are comparable in terms of biological space coverage. Indeed, the chemical space is estimated to be extremely large,^69^ and efficient sampling of this space by the training dataset is expected to have a great impact on the generalization properties of the prediction models.

We use the t-SNE algorithm^70^ on the molecule features *ψ_M_* derived from the Tanimoto kernel, as defined in Section 4.3, to visualize the resulting high-dimensional molecular space in a two-dimensional space, thus facilitating analysis. Figure 3 shows not only that LCIdb contains a much larger number of molecules than BIOSNAP, DrugBank, and BindingDB, but also that the molecules it contains are more diverse,

**Figure 3:**
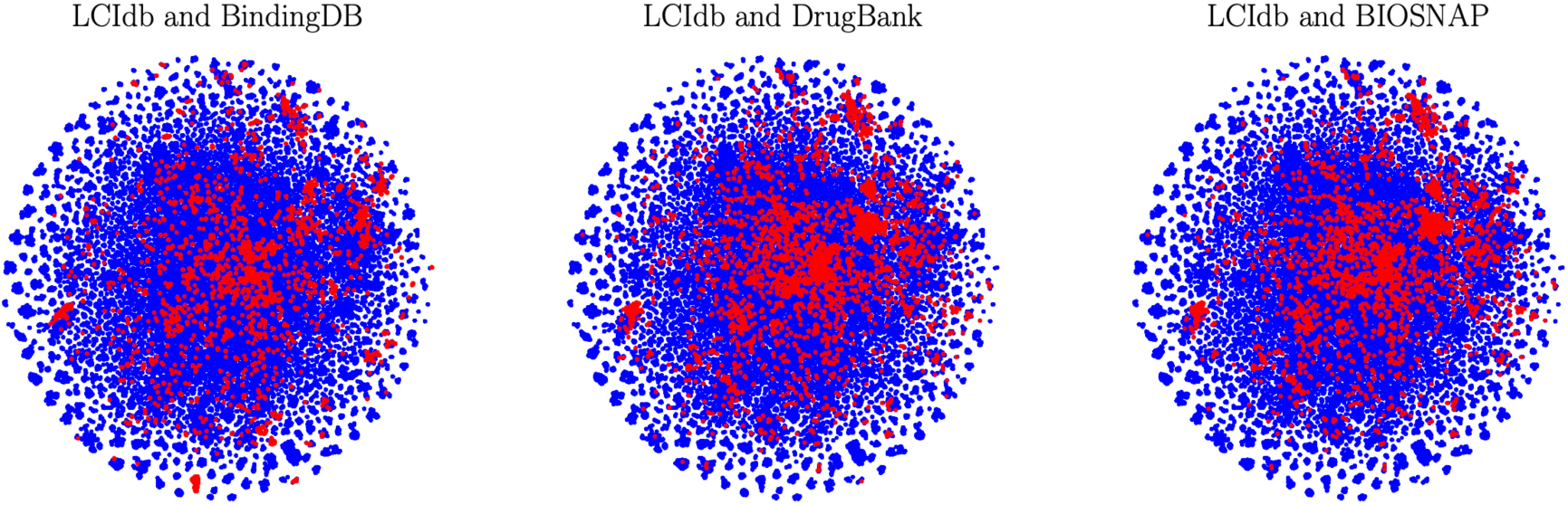
2D representation of the molecular space with the t-SNE algorithm based on molecule features. In blue: the large-sized LCIdb dataset, and in red: the medium-sized DrugBank, BIOSNAP, and BindingDB datasets.

While it is far from covering the entire vast and unknown chemical space, LCIdb spans a much larger area on the t-SNE plot, therefore providing a better sampling of this space overall. In addition, it shows that LCIdb also covers the chemical space more uniformly than the other datasets. Figure 3 also highlights that the BIOSNAP dataset was built from DrugBank, displaying similar patterns of red clusters of molecules.

We also ran the t-SNE algorithm based on Tanimoto features computed using an alternative set of molecule landmarks, and based on other molecule features (see Figure S1 of the Supporting Information). In all cases, plots confirmed the above conclusions that LCIdb presents a wider and more uniform coverage of the chemical space, underscoring their robustness.

Isigkeit *et al* ^54^ analyze the space formed by the five databases from which LCIdb originates. Specifically, they examined distributions of common drug-like features such as molecular weight, the number of aromatic bonds, the number of rotatable bonds, and predicted octanol-water partition coefficients. The authors observed that these distributions are similar across all sources. In Section 1 of the Supporting Information, we discuss plots illustrating the distribution of drugs in the LCIdb dataset, with respect to the distributions observed in the five databases from which they originate.

By contrast, the number of human proteins is comparable across all considered datasets, although not identical (see Figure 4). We also used t-SNE plots based on protein features defined in Section 4.3 to explore the coverage of the protein space by LCIdb. As shown in the resulting 2D representation presented in Figure 5, the protein space covered by LCIdb contains clusters that align with functional families of proteins. This was expected when using features calculated using the LAkernel (see Section 4.3), since proteins that share high sequence similarity usually belong to the same protein family. Thus, we can leverage this representation to discuss the diversity of proteins in our datasets. As shown in Figure 6, although LCIdb contains slightly fewer proteins than the DrugBank dataset, their coverage of the biological space is similar. BIOSNAP appears to have a lower coverage of a few protein clusters (such as protein kinases), while BindingDB focuses more on a few clusters corresponding to specific protein families.

**Figure 4:**
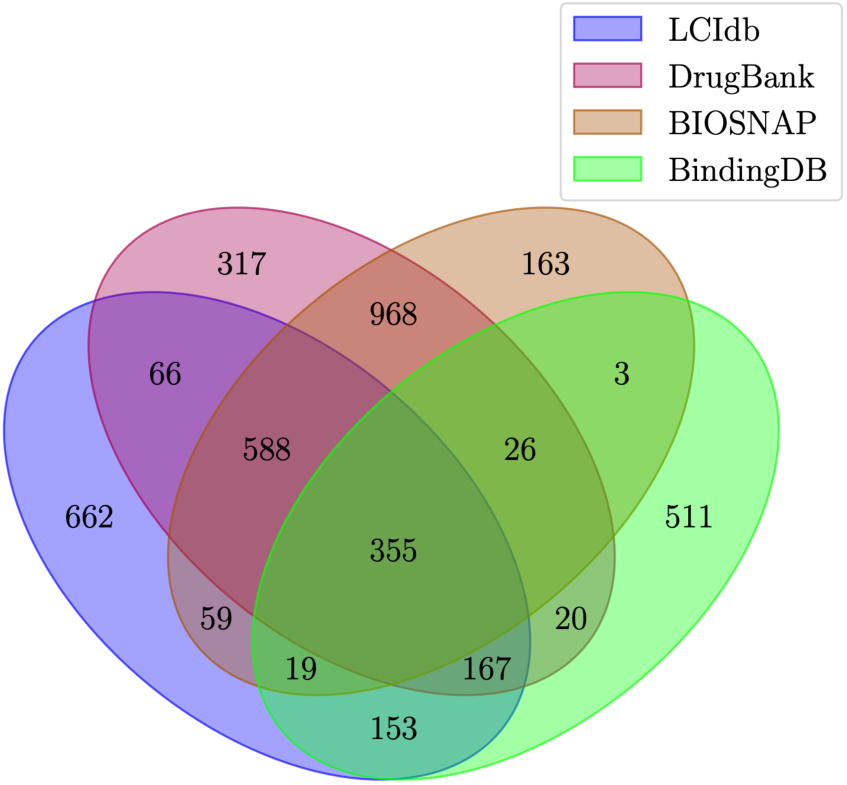
Overlap between LCIdb, DrugBank, BIOSNAP, and BindingDB datasets in terms of proteins.

**Figure 5:**
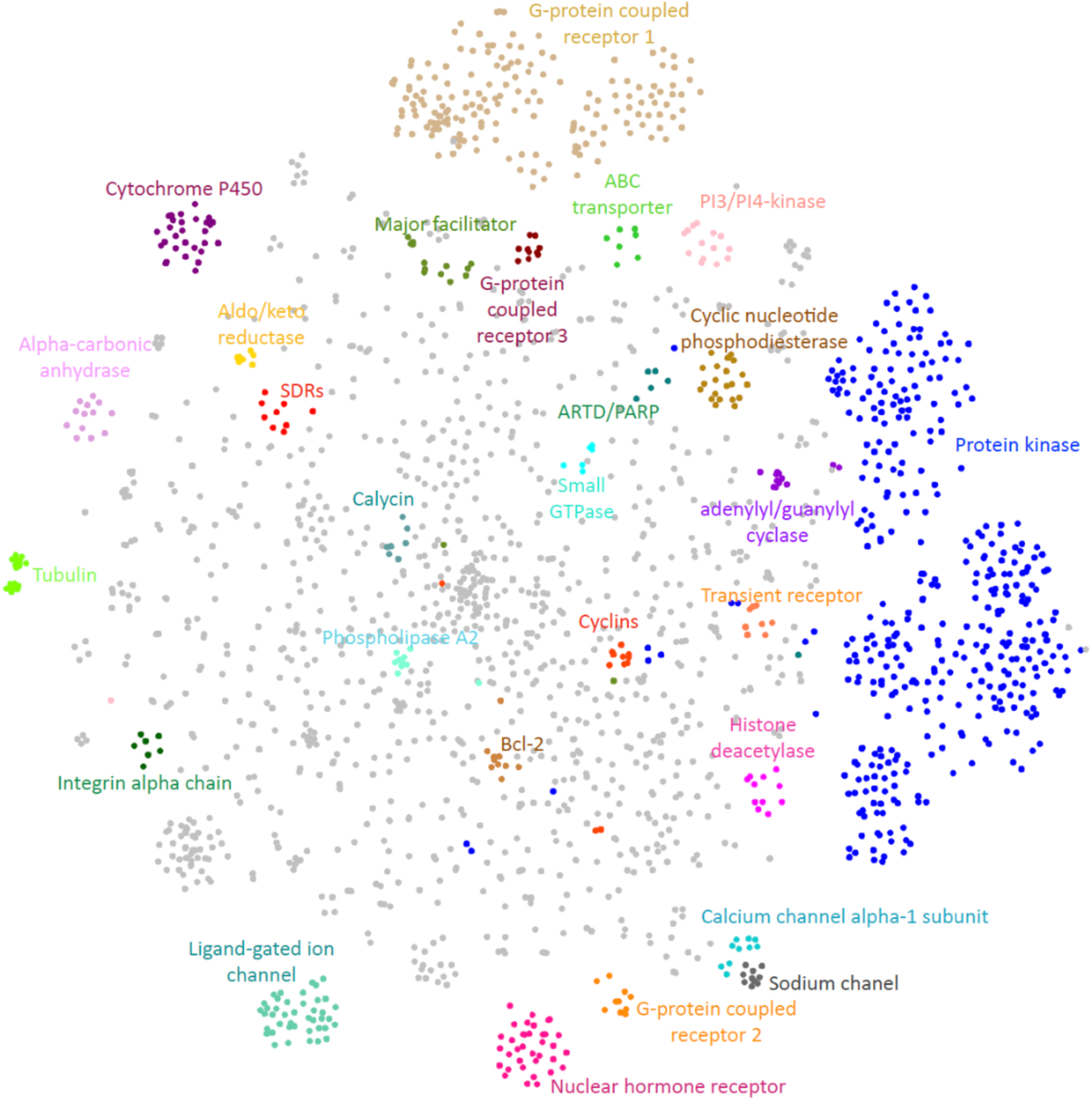
Representation of the protein space in LCIdb according to the t-SNE algorithm based on protein features derived from the LAkernel. A few protein families are labelled and coloured.

**Figure 6:**
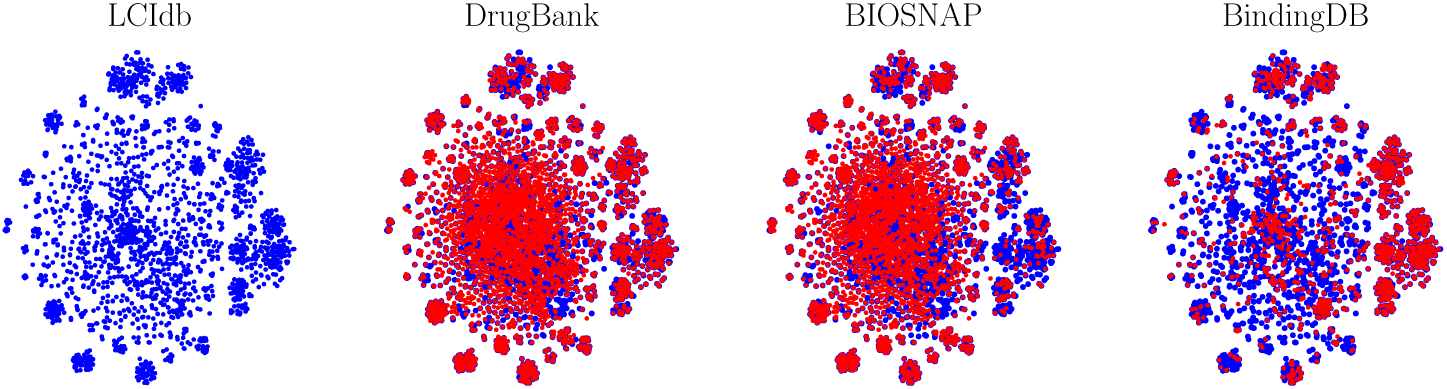
Representation of the protein space according to the t-SNE algorithm based on protein features derived from the LAkernel. In blue: LCIdb, in red: DrugBank, BIOSNAP, and BindingDB.

As detailed in Section 4.1, for BIOSNAP and LCIdb, additional datasets are derived, as suggested in various studies,^12,13,29,39,71^ as well as in Huang et al. ^20^ and Singh et al. ^49^, two papers that respectively introduced the MolTrans and ConPLex algorithms. They correspond to various scenarios in drug discovery projects: (1) the Unseen_drugs case is typical of new drugs identified in phenotypic screen and for which targets are searched to elucidate the drug’s mechanism of action; (2) the Unseen_targets case is typical of newly identified therapeutic targets against which potential drug repositioning opportunities are searched; (3) The Orphan case is typical of a new therapeutic target against which ligands (inhibitors or activators) are searched.

The composition of the corresponding datasets is provided in Table 1. In Huang et al. ^20^ and Singh et al. ^49^, only the Unseen_drugs and Unseen_targets were considered, but we added the Orphan case for LCIdb, which corresponds to the most difficult scenario.

### 5.2 Parameters set-up of the model

Due to the large number of molecules in LCIdb (see Table 1), Komet incorporates the Nyström approximation to compute molecule features using *m_M_* landmark molecules, with potentially further dimension reduction to *d_M_ ≤ m_M_* during the SVD decomposition, as presented in Section 3.2 and detailed in Section 4.3. By contrast, for proteins, we retain all the proteins in the train set as protein landmarks (*n_P_* = *m_P_* = *d_P_*), because the number of proteins in LCIdb does not lead to computational issues. It is therefore crucial to evaluate the potential impact of the *m_M_* and *d_M_* parameters on the prediction performance of Komet, the resulting gain in calculation time, and to study whether good default values can be determined. This was performed on LCIdb_Orphan and BindingDB, which are respectively large- and medium-sized datasets. LCIdb_Orphan was chosen as the large dataset for this study because it corresponds to the most difficult dataset, on which it is critical not to degrade the prediction performances. Figure 7 shows that for both datasets, we can significantly reduce the number of landmark molecules (*m_M_*) and the dimension (*d_M_*) of molecular features without losing performance, while saving time and computational resources. In particular, results on BindingDB illustrate that reducing *m_M_* from the total number of molecules (7 161) to 5 000 or 3 000 does not significantly affect precision-recall curves. In addition, for the large-sized datasets like LCIdb_Orphan, the curves corresponding to *m_M_* from 10 000 to 5 000 or 3 000 are almost superimposed, while reducing *m_M_* to 1 000 slighlty degrades the prediction performance. Figure S3 in Supplementary Information displays AUPR values for smaller values of *m_M_*, showing that further reducing the value of *m_M_* further degrades performance.

**Figure 7:**
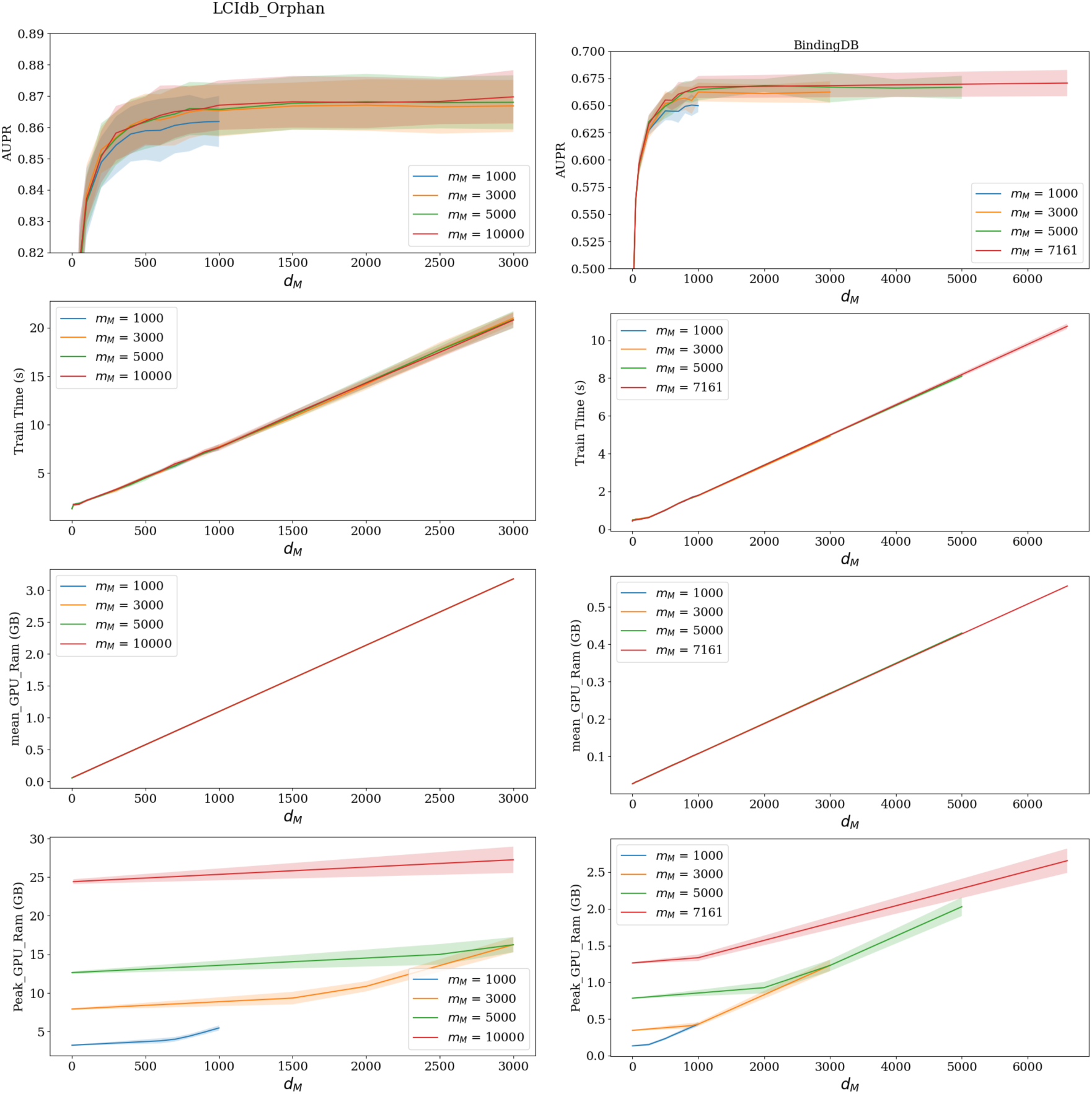
Influence of *m_M_* and *d_M_* on AUPR, computation time (in seconds) and usage and peak GPU RAM (in Gb). In each graph, the four curves correspond to four values of *m_M_*, i.e. the number of random molecules used by the Nyström approximation of the molecular kernel. Error bars are obtained by CV. Graphs on the left refer to the large-sized dataset (LCIdb_Orphan), and on the right to the medium-sized dataset (BindingDB).

Moreover, the precision-recall curves reach a plateau for *d_M_* values between 1 000 and 2 000, suggesting that we can limit the number of molecular features without a loss in performance. This observation is confirmed with the medium-size dataset BindingDB, for which a plateau is also reached for similar values of *d_M_*, particularly when no approximation was made (*n_M_* = *m_M_* = 7 161). This suggests that *d_M_* values in the range of 1 000-2 000 could be good default values for the number of features used in molecular representations. In addition, Figure 7 illustrates that, as expected, reducing *m_M_* and *d_M_* significantly reduces computational time and GPU memory usage. We finally choose *d_M_* = 1 000 and *m_M_* = 3 000 as a good compromise to design a rapid and less resource-intensive algorithm, without majorly compromising performance.

### 5.3 Impact of different molecule and protein features on Komet prediction performances

We explored the impact of molecule and protein features on the prediction performances of Komet. For molecule features, we consider the features extracted from the Tanimoto kernel between ECFP4 fingerprints, as described in Section 4.3, with the ECFP4 fingerprints themselves. This is equivalent to using the dot product between ECFP4 fingerprints, rather than the Tanimoto kernel, and no approximation (neither through the choice of a reduced set of landmark molecules nor through dimensionality reduction). Previous studies have shown that ECFP4 fingerprints perform as well as state-of-the-art fingerprint-based 3D models,^72^ and are not significantly outperformed by features learned from deep learning methods.^73^ Therefore, we also considered pre-trained Graph Neural Networks (GNNs) for the generation of molecule features. Specifically, Hu et al. ^16^ outline several pre-training strategies for GNNs using a dataset of two million molecules. These strategies include supervised learning for molecular property prediction and semi-supervised learning methods such as context prediction, mutual information maximization between local and global graph representations, encouraging similarity in representations of adjacent nodes while differentiating distant nodes, and predicting masked node and edge attributes. We use the trained models adapted by Li et al. ^74^ to calculate the molecule features and we present in Table 2 only the features giving the best results. These features correspond to a supervised learning model for molecular property prediction, combined with semi-supervised learning for context prediction.

**Table 2:**
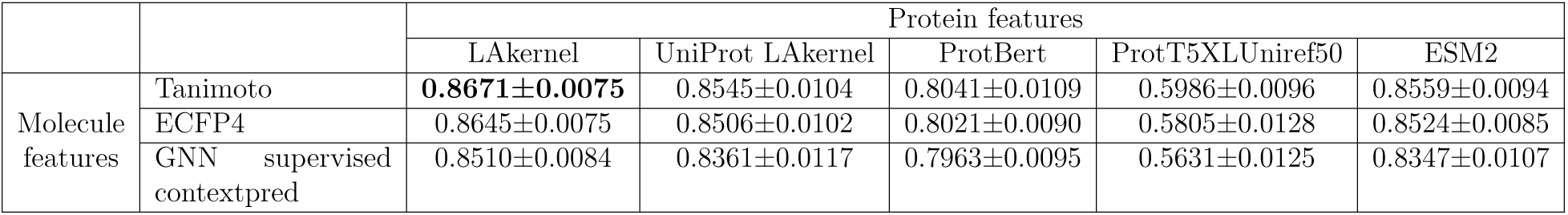
AUPR of Komet using different molecule and protein features on the LCIdb_Orphan dataset, 5-fold cross-validation. “Tanimoto” features are built from the Tanimoto kernel between ECFP4 fingerprints as described in Section 4.3, and the “GNN supervised contextpred” features are available in the DGL-LifeSci package^74^. “LAkernel” features are built from the Local Alignment kernel between proteins as described in Section 4.3. “UniProt LAkernel” features are built in the same way, but considering all human proteins from UniProt as landmarks proteins and using dimensionality reduction.

|  |  | Protein features |  |  |  |  |
| --- | --- | --- | --- | --- | --- | --- |
|  |  | LAKernel | UniProt LAKernel | ProtBert | ProtT5XLUniref50 | ESM2 |
| Molecule features | Tanimoto | **0.8671±0.0075** | 0.8545±0.0104 | 0.8041±0.0109 | 0.5986±0.0096 | 0.8559±0.0094 |
|  | ECFP4 | 0.8645±0.0075 | 0.8506±0.0102 | 0.8021±0.0090 | 0.5805±0.0128 | 0.8524±0.0085 |
|  | GNN supervised contextpred | 0.8510±0.0084 | 0.8361±0.0117 | 0.7963±0.0095 | 0.5631±0.0125 | 0.8347±0.0107 |

For proteins, we compare features extracted from the LAkernel, as described in Section 4.3, with features computed similarly, but using the 20 605 proteins of the UniProt human proteome^75^ as landmark proteins, with a dimension reduction step (*d_P_* = 1 200). In addition, we used three types of features coming from deep learning models: ESM2^25^ which is based on transformers, and ProtBert^26^ and ProtT5XLUniref50^26^ which are based on variational autoencoders trained on very large datasets of proteins.

Results are displayed in Table 2 for LCIdb_Orphan, the most challenging large-sized dataset. They show that the features proposed for Komet lead to the best prediction performance. However, replacing the molecular features built from the Tanimoto kernel between ECFP4 fingerprints with the ECFP4 fingerprints themselves barely degrades the performance. This could indicate that the molecular information lost by approximations (using a subset of landmark molecules and performing dimensionality reduction) is compensated by the Tanimoto kernel being a more appropriate kernel than the dot product. The protein features derived from the LAkernel on the 2 060 druggable proteins,^75^ i.e. human proteins for which at least one drug-like ligand is known, lead to the best prediction performances. One explanation could be that the human druggable proteins present some sequence and family bias, and do not span the whole human proteome space. As a consequence, generic features learned in deep learning approaches on very large sets of proteins from multiple species (ProtBert, ProtT5XLUniref50, ESM2), may be less appropriate for the specific problem DTI prediction in the context of drug-like molecules and human druggable proteins. This may also explain why features derived from the LAkernel computed on 20 605 human proteins also degrade the prediction performance. For this latter case, using the whole human proteome comes with the necessity of dimensionality reduction (*d_P_* = 1 200), which may also contribute to reducing the prediction performance.

As a consequence, the molecule features derived from the Tanimoto kernel on the ECFP4 fingerprints together with the protein features derived from the LAkernel on the 2 060 druggable proteins, are used in all the following prediction experiments performed with Komet. However, one should note that except for the ProtT5XLUniref50 protein features, the prediction performances of Komet remain relatively stable to molecule and protein features.

### 5.4 Comparison of the prediction performances between Komet and other algorithms

Because LCIdb is large, deep learning methods are expected to perform well on it.^76^ Therefore, we compare Komet to the recently proposed ConPLex^49^ algorithm, a deep learning approach that was shown to achieve state-of-the-art performance on medium-sized datasets.

ConPLex uses as input molecules encoded with Morgan fingerprints and proteins encoded by pre-trained Protein Language Model ProtBert.^26^ The latent space for (molecule, protein) pairs is learned through a non-linear transformation into a shared latent space. This learning phase combines a binary DTI classification phase with a contrastive divergence phase, in which the DUD-E database,^77^ comprising 102 proteins together with ligands and non-binding decoys, is used to compute a loss that minimizes the target-ligand distances (corresponding to positive DTIs) and maximizes the target-decoy distances (corresponding to negative DTIs).

We also compared Komet to MolTrans,^20^ another recent and state-of-the-art deep learning framework. MolTrans uses a representation of molecules (resp. proteins) based on frequent subsequences of the SMILES (resp. amino acid) strings, combined through a transformer module.

We finally compared Komet with simple feature-based methods. We tested a Random Forest (RF) algorithm, using concatenated molecule and protein features to create features for the (molecule, protein) pairs. We implemented the Random Forest using the scikit-learn library.^78^

#### 5.4.1 DTI prediction performances on medium-sized datasets

We first compare the performance of Komet to those of ConPLex, MolTrans and RF with concatenated features on the medium-sized datasets BIOSNAP, BindingDB and DrugBank introduced in Section 4.1. We only use the AUPR score because most negative interactions in the considered datasets are unknown interactions. The results are presented in Table 3. Note that the performance of a random predictor would correspond to an AUPR score of 0.5, except for BindingDB, where the number of negative DTIs is much greater than the number of positive DTIs. For BindingDB, the performance of a random predictor would be equal to 0.17, which explains the lower performance observed for all algorithms. We report the average and standard deviation of the area under the precision-recall curve (AUPR) in 5-fold cross-validation. Interestingly, in all cases, Komet’s AUPR performances (with *d_M_* = 1 000 and *m_M_* = 3 000) are similar to or higher than those of the two deep learning methods. This is consistent with the expectation that deep learning methods only outperform shallow learning methods when training data are abundant, due to their larger number of parameters to fit.

**Table 3:** AUPR performances of Komet, ConPLex, MolTrans and RF with concatenated protein and molecule features on medium-sized datasets BIOSNAP, BindingDB, and Drug-Bank, in 5-fold cross-validation. The ConPLex and MolTrans algorithms were re-run on these three datasets, and the resulting AUPR are very close (in fact slightly better) to those in the original paper.

| Dataset | Komet | ConPLex | MolTrans | RF with concatenated features |
| --- | --- | --- | --- | --- |
| BIOSNAP | **0.9429±0.0008** | 0.9246±0.0037 | 0.8989±0.0048 | 0.9121±0.0032 |
| Unseen_drugs | **0.8979±0.0051** | 0.8763±0.0072 | 0.8547±0.0045 | 0.8620±0.0100 |
| Unseen_targets | **0.8754±0.0099** | 0.8641±0.0100 | 0.7058±0.0273 | 0.8127±0.0116 |
| BindingDB | 0.6598±0.0074 | **0.6765±0.0178** | 0.6196±0.0150 | 0.6454±0.0075 |
| DrugBank | **0.9400±0.0030** | 0.8961±0.0070 | 0.8068±0.0100 | 0.8018±0.0086 |

In the Unseen_drugs and Unseen_targets scenarios on BIOSNAP, as expected, the AUPR performances decrease for all algorithms but remain high, except for MolTrans which overall tends to display lower performances than the two other algorithms.

#### 5.4.2 DTI prediction performances on large-sized datasets

Then, we trained Komet, ConPlex, MolTrans and RF with concatenated features on the four large-sized LCIdb-derived datasets. The results demonstrate that Komet achieves state-of-the-art prediction performance in all cases (see Table 4) at a much lower cost in terms of training time (see Table 5).

**Table 4:** Comparison of AUPR scores on large-sized datasets, in 5-fold cross-validation.

| Dataset | Komet | ConPLex | MolTrans | RF with concatenated features |
| --- | --- | --- | --- | --- |
| LCIdb | **$0.9925 \pm 0.0004$** | $0.9783 \pm 0.0008$ | $0.9721 \pm 0.0011$ | $0.9865 \pm 0.0002$ |
| Unseen_drugs | **$0.9944 \pm 0.0003$** | $0.9831 \pm 0.0009$ | $0.9710 \pm 0.0004$ | $0.9829 \pm 0.0006$ |
| Unseen_targets | **$0.8952 \pm 0.0186$** | $0.8780 \pm 0.0223$ | $0.5987 \pm 0.0131$ | $0.6886 \pm 0.0232$ |
| Orphan | **$0.8671 \pm 0.0075$** | $0.8175 \pm 0.0130$ | $0.5455 \pm 0.0004$ | $0.5961 \pm 0.0070$ |

**Table 5:**
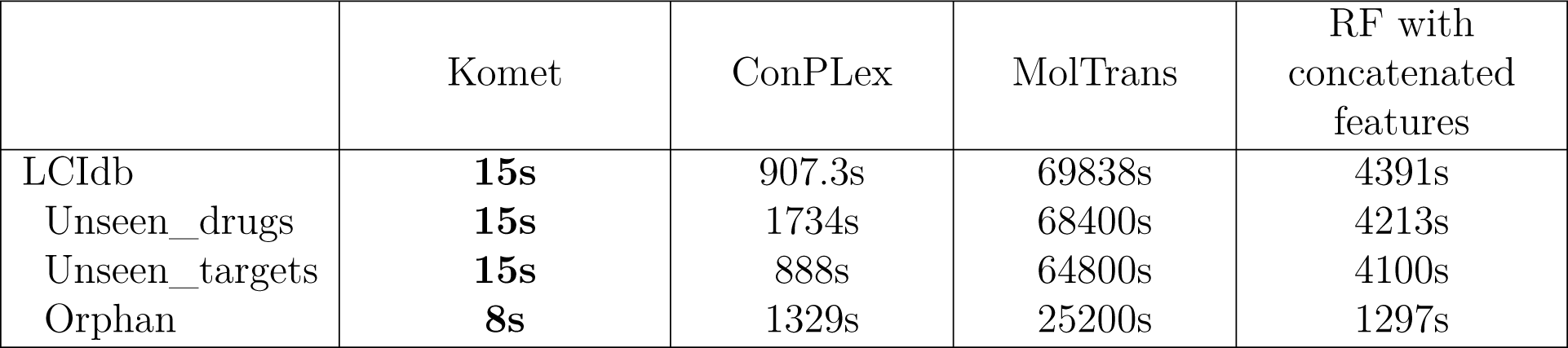
Comparison of training time for the considered algorithms.

Overall, the performance of Komet is consistently high, with AUPR scores above 0.9 in most cases. Because the number of molecules is still very large in the LCIdb Unseen_drugs dataset, thus covering a broad chemical space, the performance remains excellent, although molecules in the Test set are absent in the Train set. In LCIdb Unseen_targets and LCIdb_Orphan, where the proteins in the Test set are absent in the Train set, the performances are slightly lower but remain high. The ConPLex algorithm also displays high performances (although lower than those of Komet) in all cases, while MolTrans and RF appear less efficient in these more stringent scenarios. However, Komet is much faster. In addition, although using the same molecule and protein features, Komet is much faster than RF on large datasets like LCIdb, and performs much better in scenarios where fewer data is available (unseen targets or orphan). Our interpretation is that the key aspect in Komet (besides is scalability) is the use of the Kronecker product of the protein and molecular spaces to encode (molecule, protein) pairs.

We conducted a comparison using various performance measures, and the outcomes consistently align with the above results. For these additional insights, please refer to Section 3 of the Supporting Information for details.

#### 5.4.3 Validation on DrugBank (Ext) as external dataset

In the above sections, the performances of the algorithms are compared in 5-fold cross-validation for all datasets. To better assess the generalization properties of the algorithms, we used as an external dataset the DrugBank (Ext) introduced in Section 4.1.

The prediction performance of the three considered algorithms on DrugBank (Ext), when trained on BindingDB or on LCIdb, are reported in Table 6, from which two conclusions can be drawn. First, all ML algorithms perform better when trained on LCIdb compared to BindingDB. This improvement is attributed to LCIdb’s more large coverage of both chemical and protein spaces. Indeed, according to Figure 3, the molecule space covered by LCIdb globally includes that covered by DrugBank, but this does not appear to be the case for the BindingDB dataset. Similarly, according to Figure 5, the protein space of LCIdb globally covers that of DrugBank, whereas the protein space of BindingDB does not seem to cover that of DrugBank.

**Table 6:** AUPR performance for the considered algorithms trained on BibndingDB and LCIdb

| Training set \ Algorithm | Komet | ConPLex | MolTrans |
| --- | --- | --- | --- |
| LCIdb | **0.848** | 0.822 | 0.558 |
| BindingDB | **0.659** | 0.611 | 0.503 |

Second, Komet always outperforms the two deep learning algorithms. Overall, Komet trained on LCIdb displays the best generalization performances on DrugBank (Ext).

### 5.5 Large-scale predictions in the chemical space: application to large-step scaffold hopping problems

The need for scaffold hopping is a recurrent problem in drug design. It refers to situations where a global structure of a hit molecule against a protein target is not suitable, because of unacceptable toxicity, poor ADME profiles, lack of specificity or selectivity, or protection by a patent that restricts its development. The goal is then to identify new molecules with similar bioactivity that the initial hit, i.e. bind to the same protein pocket with similar binding modes. This demanding task corresponds to an important challenge in drug discovery.^3^ Depending on the cases, one will need to search for new molecules with various degrees of similarity with respect to the hit, which led to distinguishing small-, medium-, and large-step scaffold hopping cases, referring to how far from the hit we need to jump in the chemical space.^79^ Various strategies can help design new molecules from the initial hit, including subtle changes made to the connecting fragments or substituents of the hit, heterocycles ring opening or closure, swapping of carbons and heteroatoms in heterocycles, or designing new molecules thanks to topology-based (3D) approaches or various computational approaches.^79,80^ Although various examples of successful scaffold hopping cases have been reported, these types of problems remain difficult to solve without the aid of computational methods,^79^ and new concepts are particularly required to help solve the most difficult cases, i.e. large-step scaffold hopping cases.

Solving large-step scaffold hopping problems is related to the topic of the present paper because it requires DTI prediction at large scales in the chemical space. Indeed, the model should display good prediction performances broadly in this space, so that it may be reliable far from the initial hit. Therefore, although Komet was not dedicated to scaffold hopping, it can be trained on the very large LCIdb dataset in order to best cover the chemical space, which led us to evaluate its interest as a tool for solving such problems.

In a previous study, we proposed the *ℒℌ* benchmark to assess the performance of computational methods to solve large-step scaffold hopping problems.^44^ As detailed in Section 4.6, the *ℒℌ* benchmark is a high-quality dataset comprised 144 non-redundant pairs of highly dissimilar molecules that bind to the same protein pocket with similar binding modes. These pairs were identified from PDBbind, and are well-characterized examples of “true” large-step scaffold hopping cases for a panel of 69 proteins belonging to various diverse families. On this benchmark, computational methods are evaluated as follows: given one molecule of a pair (the known active), the objective is to rank the other (the unknown active) among 499 decoy molecules. The lower this rank, the better the prediction performance. This allows to simulate the requirements for real-case applications, where only the best-ranked molecules would be experimentally tested. Since either molecule of the pair can be assigned as the known active, this leads to 288 scaffold hopping cases to solve.

In Figure 8, we compare the performance of the Komet and ConPLex algorithms trained on LCIdb or BindingDB, using Cumulative Histogram Curves (CHC). This criterion illustrates the frequency of cases where the method ranked the unknown active molecule below a given rank. Table 7 provides the Area Under the Curve (AUC) of CHC curves, offering a quantitative comparison of methods, along with the proportion of cases where the unknown active was retrieved within the top 1% and 5% of best-ranked molecules. These metrics also serve as indicators of the success rate of the methods. We re-computed the results obtained by the Kronecker kernel with an SVM calculated with the scikit-learn toolbox, using the same kernels as in Komet, but trained on the DrugBank dataset, in order to evaluate the impact of the chemical space coverage provided by the training set. These results align with those of the original paper by Pinel et al. ^44^.

**Figure 8:**
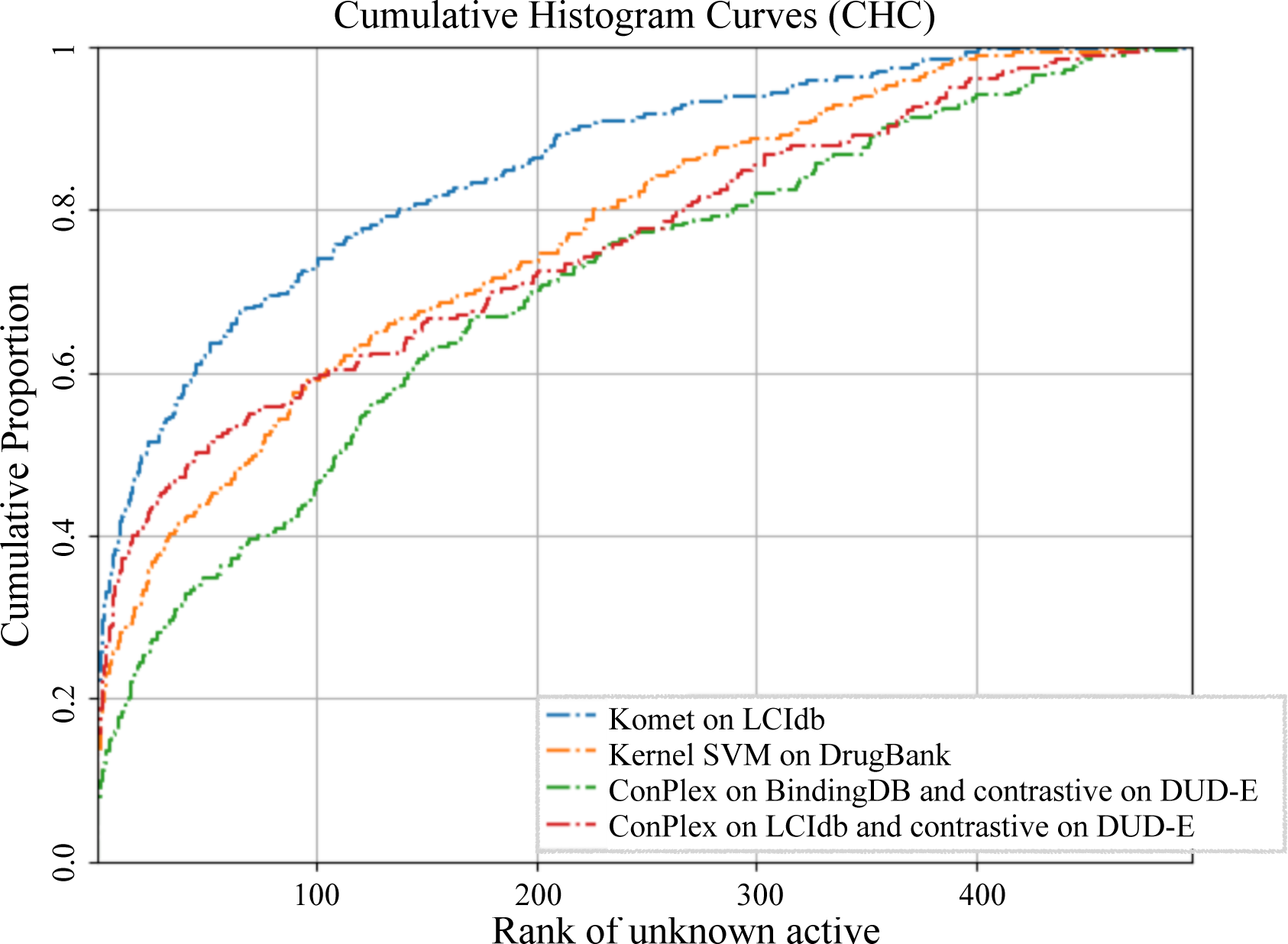
Cumulative Histogram Curves of the considered algorithm, measuring the cumulative proportion of cases the unknown active is retrieved below a given rank.

**Table 7:** Prediction performances on the *ℒℌ* benchmark.

| Dataset | Komet<br>on<br>LCIdb | Kernel SVM<br>on DrugBank | ConPlex on BindingDB<br>and contrastive on DUD-E | ConPlex on LCIdb<br>and contrastive on<br>DUD-E |
| --- | --- | --- | --- | --- |
| ROC-AUC | **0.85** | 0.77 | 0.70 | 0.75 |
| Top 1% | **32%** | 22% | 12% | 24% |
| Top 5 % | **52%** | 36% | 26% | 43% |

As shown in Figure 8 and Table 7, the performances of Komet and ConPlex improve when trained on LCIdb over those observed when trained on BindingDB, while the kernel SVM trained on DrugBank displays performances that are intermediates with those of ConPlex on the two training datasets. This is consistent with a broader chemical diversity in LCIdb, offering a better coverage of active molecules in *ℒℌ* than BindingDB and DrugBank. Indeed, we used the t-SNE algorithm to visualize that LCIdb uniformly spans the entire space of active molecules in *ℒℌ*, which is not the case for the DrugBank and the BindingDB datasets (see Figure 9)

**Figure 9:**
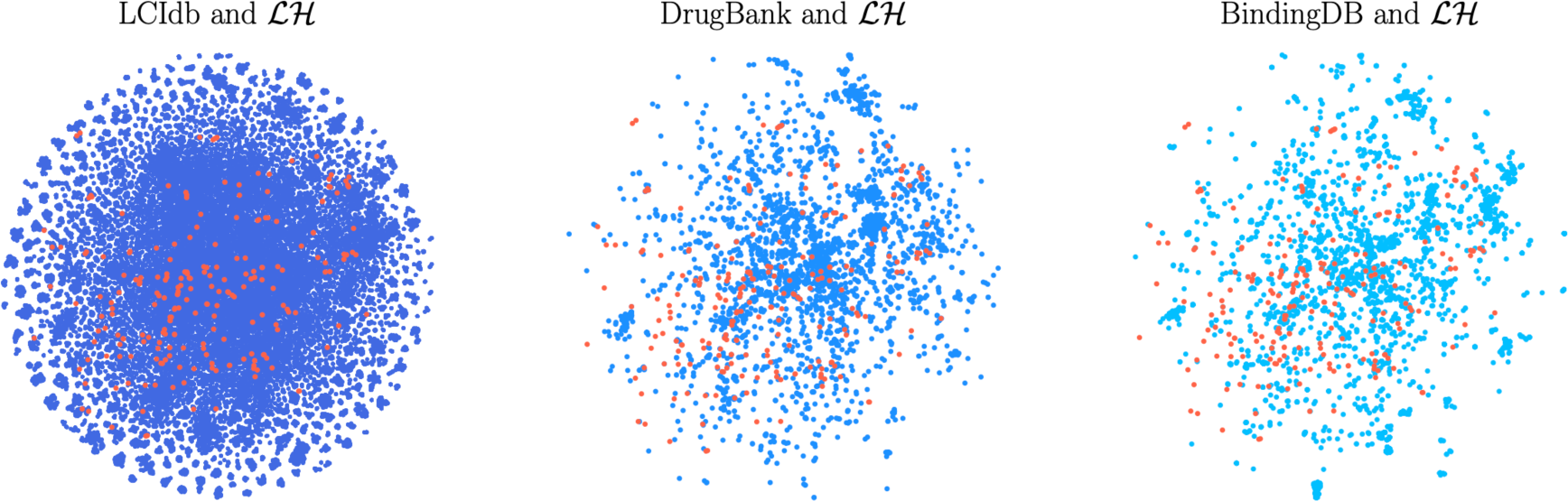
t-SNE on molecule features. In blue and from left to right: LCIdb, DrugBank and BindingDB, in orange: active molecules of *ℒℌ*.

It is somewhat puzzling that, in this specific study, ConPLex does not outperform Komet when both are trained on the same dataset. Indeed, we chose ConPLex because it incorporates a contrastive learning step based on the DUD-E database, which should help separate the unknown positive from the decoys in *ℒℌ*. One explanation may be that DUD-E presents a hidden bias that was shown to mislead the performance of deep learning algorithms.^81^ The use of an unbiased database for contrastive learning may improve the performance of ConPLex on the *ℒℌ* benchmark.

In addition, the fact that Komet outperforms ConPLex may illustrate that tensor product-derived features for the (molecule, protein) pairs used in Komet better capture interaction determinants than the features learnt by the deep learning algorithm in ConPLex. It is consistent with the fact that the SVM with the tensor product kernel, although trained on DrugBank, also performs better than ConPlex trained on LCIdb.

Overall, Komet trained on LCIdb successfully ranks the unknown active in the top 5% in 50% of cases. This performance surpasses those of all ligand-based methods tested in the original paper by Pinel et al. ^44^, the best of which, involving 3D pharmacophore descriptors, ranked the unknown active in the top 5% in 20% of cases. However, these results remain halftone and still leave a lot of space for improvement. We do not claim that Komet should replace other approaches, in particular 3D-based approaches, but it constitutes another string to the bow of available methods for addressing these challenging problems.

## 6 Discussion

An important contribution of the present work resides in providing the LCIdb DTI dataset, which appears much larger than most public datasets used in the recent literature. A key feature of this dataset is a wider and more uniform coverage of the molecular space. A recurrent problem when building DTI datasets for training ML algorithms is that negative interactions are usually not reported. One way to circumvent this problem is to use reference databases that provide quantitative bioactivity measurements and choose threshold values to define positive and negative interactions. In previous studies,^20,49^ other authors chose a common and rather low threshold value of 30 nM for both types of DTIs, leading to a modest number of positive (9 166) and three times more negative DTIs (23 435), as shown in Table 1. The notion of positive and negative DTIs is not absolute, because bioactivities are continuous, and threshold values are somewhat arbitrary. In the present paper, we chose distinct thresholds for positive and negative interactions, respectively under 100 nM (10^-7^M) and above 100*µ*M (10^-4^M). This leads to a limited number of known negative DTIs in the dataset (7 965) compared to known positives (396 798). Overall, our goal was to limit the potential false negative DTIs and the bias towards well-studied molecules and proteins. Therefore, true negative DTIs were completed by randomly chosen DTIs according to the algorithm in Najm et al. ^50^, while excluding all DTIs with activities falling in the 10^-4^ - 10^-7^M range. However, we are aware that using a lower threshold value for the negative DTIs in LCIdb would have allowed us to select a high number of DTIs considered as known negative. Furthermore, we can easily create datasets with different thresholds for defining positive/negative interactions by simply adjusting these thresholds and rerunning the code. For example, if we are mainly interested in off-target predictions rather than in the identification of primary targets, we can use a higher concentration threshold of 10 *µ*M to define positive DTIs. To illustrate this, we created the LCIdb_Orphan_10*µ*M_threshold dataset like LDIdb_Orphan, but using a threshold of 10 *µ*M rather than 100nM. Training Komet is still fast, even with this larger training set (1 037 934 positive DTI, 508 353 molecules, and 2 970 proteins). We evaluated the considered algorithms on this new dataset, on the most challenging scenario (Orphan), using 5-fold cross-validation. The results, provided in Section 4 of the Supplementary Information, demonstrate that Komet still outperforms the other considered algorithms.

The Komet algorithm has two parameters, *m_M_* (number of landmark molecules) and *d_M_* (dimension of molecular feature vectors). We were able to define good default values for these parameters (*d_M_* = 1 000 and *m_M_* = 3 000), significantly reducing the computational time and memory requirements. Interestingly, computational resources will not increase drastically if the size of the train set increases (i.e. if new DTIs are added), as can be judged from Figure 7. The number of proteins in LCIdb (2 060) is much smaller than the number of molecules (271 180), so we did not need to use the Nyström approximation or perform reduction of dimension to compute protein feature vectors. This means that the dimension of protein feature vectors is *n_P_* = *m_P_* = *d_P_*. However, should Komet be trained on other datasets containing many more proteins, smaller values of the *m_P_* and *d_P_* could be used, because in the implementation of Komet, computation of protein and molecule features is treated similarly.

We also showed that the performance of the algorithm was robust for the choice of the landmark molecules and the molecule and protein features, although learned features tended to decrease the performance, as shown in Table 2.

Importantly, Komet belongs to the family of shallow ML algorithms and proved to outperform ConPLex and MolTrans, two recently proposed deep learning algorithms, at a much lower computational cost.

One explanation for the good performance of Komet could be that features for the (molecule, protein) pairs derived by Komet in Step 2, simply based on the Kronecker product, may better capture determinants of the interaction than the concatenated features used in the RF algorithm, or the combined features learned in the considered deep learning algorithms. Interestingly, the choice of fixed or learnt molecular features did not significantly impact performance (see Table 2). In contrast, learnt protein features tended to decrease the performances.

In addition, by using the algebraic properties of the tensor product, an efficient implementation allows the use of a quasi-Newton optimization algorithm, in full batch. Thus, Komet is easier and faster to optimize than DL models, which often require more complex and time-consuming training procedures. The choice of the tensor product not only provides excellent scalability to Komet but also enables it to leverage the rich information present in the very large LCIdb dataset effectively. This scalability ensures that Komet can handle extensive datasets without a significant increase in computational resources, making it a robust choice for large-scale DTI predictions.

Furthermore, our study focuses on DTI prediction in the human druggable space of proteins, because our goal is to propose a tool for drug discovery projects. The dimension of this space is modest, as illustrated by the number of proteins in LCIdb (2 060), with respect to that of the human proteome (above 20 000, but expected to be in the order of 90 000 when including splicing variants). Therefore, the druggable human proteins may present some sequence bias, and the protein features used in ConPLex and MolTrans and learned based on a much wider space of proteins may not be optimal for the DTI prediction of the problem at hand. This is consistent with the results in Table 2, showing that learned features did not improve the performances of Komet.

Overall, Komet proved to display state-of-the-art performances on various prediction scenarios that require large-scale prediction, such as de-orphanization or scaffold hopping problems.

One limitation of Komet is that the protein and molecule features and kernels are somewhat rough. Indeed, when the goal is to train models with broad applicability domains, features for proteins and molecules can only be computed based on simple representations such as primary sequence and 2D chemical structure, respectively. Indeed, richer molecule features encoding protonation state, 3D conformation of the bound ligand, 3D shape or 3D pharmacophore information, or type of interactions with the protein can only be computed is not feasible at large scales. In particular, some of these features would rely on 3D information, which is not available for the 271 180 DTIs in LCIdb. The same situation holds for proteins, and richer protein features encoding the protein binding pockets cannot be derived at this scale. This also has as a consequence that Komet can only predict whether a molecule is expected to bind a protein, but not where. In particular, it does not provide any information about the binding pocket to which the molecule may bind when the protein has several.

Local models trained on smaller datasets focusing on particular protein families (kinases, GPCRs, ion channels, etc…), may leverage richer information available in these families into more sophisticated features. For example, in the case of GPCRs, it has been shown that an SVM algorithm with specific kernels based on amino acids defining the binding pockets or on the hierarchical structure of this family, displays better prediction performances than the LAkernel used in the present study.^28^ Therefore, in specific protein families, such local models could outperform Komet in its present version, but these models would not be applicable at large scales, such as on LCIdb. Note however that Komet is a versatile algorithm that could also be trained on a smaller dataset of DTIs involving specific protein families, in which richer features could be computed and used as input, leading to a local version of Komet optimized for this family.

Komet uses a multi-task approach, that is to say, makes use of information about interactions that involve neither the query protein nor the query ligand. In order to showcase the benefit of such an approach in the case of protein de-orphanization, we compared Komet to a baseline model trained only on DTIs involving the nearest protein (NN) in the dataset (as the protein itself, being orphan, cannot provide any positive training DTI to a single-task model). To make a fair comparison, we considered a linear SVM using the same molecular features as Komet, so that the NN model’s algorithm and molecular encodings are comparable to those of Komet. We computed the AUPR for both models in different settings: on all proteins in LCIdb„ and on several subsets of proteins in LCIdb: GPCR receptor proteins, kinases, proteins whose NN is close (for which the LA kernel with the NN is higher than 0.75), and proteins whose NN is far (for which the LA kernel with the NN is lower than 0.25). Details about these experiments are provided in Supplementary Information. As shown in Table S4, in all cases, Komet outperforms the NN model. This illustrates the goal that was pursued: propose a pipeline that displays on average good performances at large scales, in a broad range of settings. However, in a few cases, we found that the NN model performed better than Komet. These cases correspond to query proteins with a small number of known ligands (that form the test set) and whose NN have many known ligands, or to query proteins that have several ligands in common with those of their NN. For example, the GPBAR1 protein, belonging to the GPCR1 family, has 3 known ligands, while its NN S1PR4, a protein from the same family, has 48 ligands. In this case, the AUPR for the NN model is 1, versus 0.64 for Komet. Another example, the ERCC5 protein, from the XPG/RAD2 endonuclease family, has 21 ligands, and its nearest neighbour is FEN1, a protein from the same family, shares 17 ligands with ERCC5. In this case, the AUPR for the NN model is 1, versus 0.58 for Komet. The NN model also performs better than Komet in cases where the NN has many known ligands, while the query protein has very few. For example, the GPBAR1 protein, from the GPCR1 family, has 3 known ligands, while S1PR4, its NN belongs to the same family and has 48 ligands. Here, the AUPR for the NN model is 1, versus 0.64 for Komet.

As mentioned above, local models with specific features tailored to a particular protein family could improve the performances in Table S4. But these improvements would apply both to NN models and to a local model of Komet fueled with these specific features.

## Data and Software Availability

We use a server with 2 CPUs and 1 NVIDIA A40 GPU with 48 GB of memory. We provide a Python implementation of Komet and the code used to build LCIdb at https://komet.readthedocs.io. We provide the LCIdb itself and other files at https://zenodo.org/records/10731712 and https://github.com/Guichaoua/komet/tree/main/data.

## Supporting Information

The Supporting Information File offers an analysis of the molecule space coverage of the various datasets, as shown in Figures S1 and S2. It includes a study on the impact of significantly reducing the number of molecule landmarks, detailed in Figure S3. The file also provides alternative data splitting methods (Train/Validation/Test) and several metrics to compare prediction performances, presented in Table S1. Additionally, the Supporting Information contains a study on performance using another filtered dataset to examine off-target predictions. It also provides mathematical details of Section 4.4. Finally, it includes a comparison between the Nearest Neighbour single-task approach and the multi-task approach in Komet.

