## Supporting Information for "Drug-Target Interactions Prediction at Scale: the Komet Algorithm with the LCIdb Dataset"


This Supporting Information provides a complementary analysis of various aspects of our study. It includes an analysis of the molecule space coverage across different datasets, as shown in Figures S1 and S2. The impact of significantly reducing the number of molecule landmarks is explored in detail in Figure S3. Alternative data splitting methods (Train/Validation/Test) are discussed, and several metrics for comparing prediction performances are presented in Table S1. Additionally, this file contains a study on performance using a filtered dataset to examine off-target predictions. Mathematical details related to Section 4.4 are provided for deeper insights into our methodology. Finally, a comparison between the Nearest Neighbor single-task approach and the multi-task approach in Komet is presented.

### 1 Content analysis for the considered databases and for LCIdb

This section shows cases of a 2D visualization of the chemical space covered by various datasets considered in the paper, using the t-SNE algorithm on various molecule features.

Figure S1 shows the distribution of the molecules in LCIdb and in the five databases from which it was built,[1] based on a 2D visualization of the chemical space, according to the t-SNE algorithm on various molecule features. It highlights a significant contribution from the ChEMBL and PubChem databases, enhanced mainly by data from Probes&Drugs.

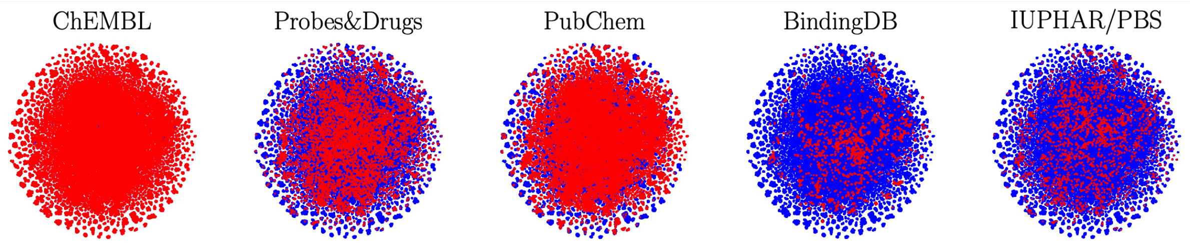


Figure S1: t-SNE on molecule features. In blue: large-sized benchmark LCIdb, in red: 5 databases from which the initial dataset[1] is extracted.

We note the importance of molecules present in ChEMBL in the LCIdb database. To evaluate the contribution of ChEMBL to LCIdb we applied to the ChEMBL database the same filters as those used to build LCIdb (see Section 4.2 and Figure 2). We obtained a dataset (available on Zenodo at `https://zenodo.org/records/10731712`) containing 1 818 proteins (compared to 2 060 in LCIdb), 267 377 molecules (compared to 271 180 in LCIdb), 386 652 positive DTIs (compared to 396 798 in LCIdb), and 7 373 negative DTIs (compared to 8,041 in LCIdb). This shows that 98.6% of the molecules and 96% of the DTIs of LCIdb come from ChEMBL. However, when using only ChEMBL, we lose more than 10% of the proteins of LCIdb, which is not negligible. The loss of the corresponding proteins can pose a problem, particularly in application where the goal would be to predict the protein interaction profile of a molecule. Furthermore, by limiting the data to only ChEMBL, we lose around 8% of negative DTIs, which are already very few compared to the number of positive DTIs.

Finally, combining ChEMBL with other databases allows us to increase our confidence in these interactions and filter out 2 180 DTIs found positive in ChEMBL but not elsewhere.

We evaluated the performance of Komet on this slightly smaller dataset (with respect to LCIdb) on the external DrugBank (Ext) dataset. When training on this new dataset, we obtained an AUPR of 0.8435 versus 0.8486 for LCIdb, which is only slightly better. However, the important difference between LCIdband the ChEMBL-derived dataset is that the former is better cleaned and more complete, providing prediction for 10% additional proteins.

Figure S2 shows the t-SNE visualizations of the molecular space for various considered datasets, based on Tanimoto features (as in Figure 2 of the article) for one choice of 3000 landmark molecules, for another choice of 3000 landmark molecules, and ECFP4 features. It confirms that LCIdb offers broader and more uniform coverage of the chemical space than BindingDB, DrugBank, or BIOSNAP.

### 2 Study of the impact of reducing the number of landmark molecules $m_M$

Figure S3 shows the Area Under the Precision-Recall curve (AUPR) as a function of the number of landmark molecules ($m_M$), with $d_M = m_M$, as $d_M$ cannot exceed $m_M$, for small values ranging from 1 to 1000. This analysis demonstrates that reducing $m_M$ to 300 causes a slight decrease in performance. Overall, given that the computational cost corresponding to $m_M = 3000$ and $d_M = 1000$ is cheap (in terms of training time, peak RAM and mean GPU, as shown in Figure 7 of the article), we argue that these values are a good choice and use them throughout our study.

### 3 Data Splitting and Performance Evaluation

Huang et al.[2] and Singh et al.[3] proposed to train and compare the performance of various

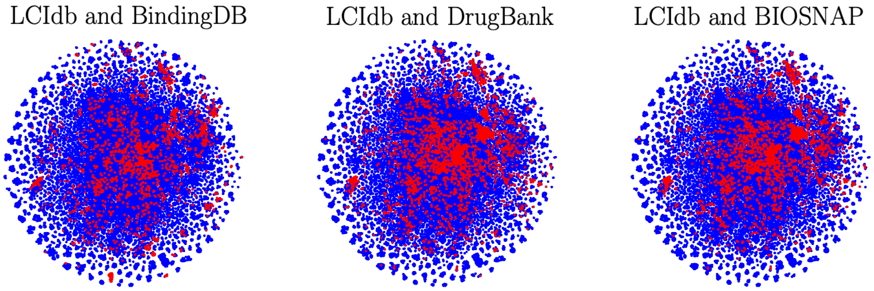


a) t-SNE visualisation on Tanimoto features

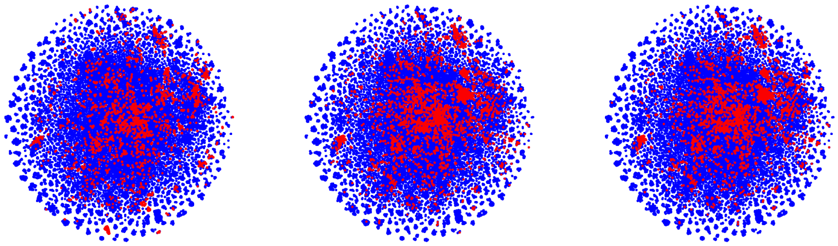

b) t-SNE visualisation on Tanimoto features, with another choice of the 3000 landmark molecules

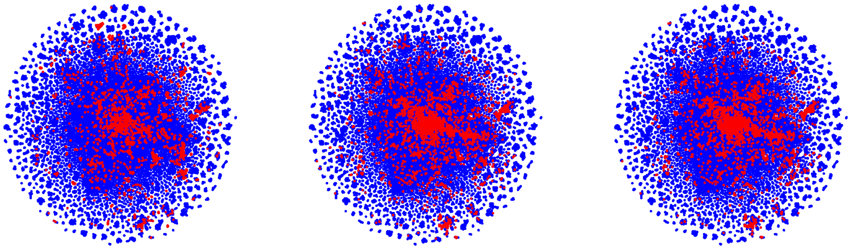

c) t-SNE visualisation on ECFP4 features

Figure S2: 2D representation of the molecular space, based on the t-SNE algorithm on molecule features. In blue: for 3000 landmark molecules in the large-sized LCIdb dataset, and in red: medium-scale DrugBank, BIOSNAP, and BindingDB datasets.

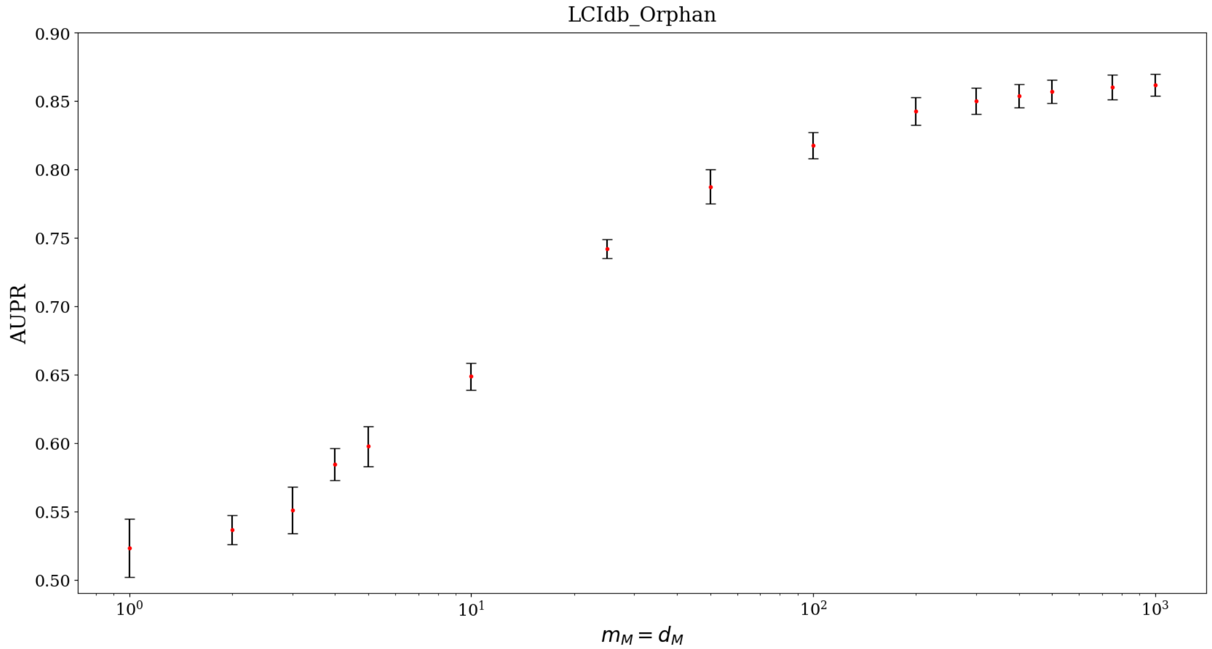


Figure S3: Influence of $m_M = d_M$ on AUPR on the large-sized dataset LCIdb_Orphan. Error bar are obtained by cross-validation.

DTI prediction algorithms based on splitting the datasets in training (Train), validation (Val), and test (Test) sets. In this section, we first present these Train/Val/Test sets and then compare the prediction performances using several metrics.

#### 3.1 Presentation of Train/Val/Test Sets

We first re-use the publicly available medium-sized datasets of Huang et al.[2] and Singh et al.[3]. Using the same scheme, we randomly split our large-sized dataset into training (Train), validation (Val), and test (Test) sets using a 7:1:2 ratio for all different prediction scenarios: LCIdb, LCIdb_Unseen_drug, LCIdb_Unseen_protein, and LCIdb_Orphan (unseen molecule and protein) scenarios. This method ensures a fair comparison of the performance of various DTI prediction algorithms. Table S1 provides the number of positive and negative interactions across the Train, Val, and Test sets in these datasets. All datasets of this section can be found at `https://zenodo.org/records/10731713`.

Table S1: Full specification of the Train/Val/Test sets for all datasets. DrugBank (Ext) is only used as an external validation dataset when algorithms are trained on BindingDB or LCIdb. Therefore, no Train, Val, or Test sets were built for DrugBank (Ext)

| Datasets | #Train | #Val | #Test |
|---|---|---|---|
| BIOSNAP | 9,670/9,568 | 1,396/1,352 | 2,770/2,727 |
| Unseen_drugs | 9,535/9,616 | 1,383/1,353 | 2,918/2,675 |
| Unseen_targets | 9,876/9,499 | 1,382/1,386 | 2,578/2,762 |
| BindingDB | 6,334/6,334 | 927/5,717 | 1,905/11,384 |
| DrugBank | 10,972/10,972 | 1,098/1,098 | 1,645/1,645 |
| DrugBank (Ext) | - | - | 10,838/10,838 |
| LCIdb | 161,015/161,015 | 32,204/32,204 | 48,304/48,304 |
| Unseen_drugs | 156,942/156,942 | 32,326/32,326 | 56,328/56,328 |
| Unseen_targets | 154,683/161,015 | 32,349/32,349 | 60,822/60,822 |
| Orphan | 59,132/59,132 | 10,145/10,145 | 22,503/22,503 |

#### 3.2 Several metrics to compare prediction performances

Table S2 presents various metrics for comparing prediction performances on the four LCIdb-datasets. While ConPlex has better accuracy in two cases, overall, Komet outperforms the other algorithms in most cases according to AUPR, ROC-AUC and Accuracy prediction performances, supporting the main conclusions in the paper.

Table S2: AUPR, ROC-AUC and Accuracy prediction performances

| | Komet | | | ConPLex | | | MolTrans | |
|---|---|---|---|---|---|---|---|---|
| | AUPR | ROC-AUC | Accuracy | AUPR | ROC-AUC | Accuracy | AUPR | ROC-AUC |
| LCIdb | **0.990** | **0.990** | **0.966** | 0.970 | 0.971 | 0.917 | 0.967 | 0.970 |
| Unseen_drugs | **0.994** | **0.994** | **0.976** | 0.980 | 0.977 | 0.934 | 0.968 | 0.969 |
| Unseen_targets | **0.915** | **0.896** | 0.714 | 0.893 | 0.874 | **0.763** | 0.591 | 0.584 |
| Orphan | **0.896** | **0.879** | 0.682 | 0.845 | 0.834 | **0.689** | 0.552 | 0.536 |

### 4 Impact of activity thresholds chosen for the specific problem of off-target prediction

We can easily create datasets with different choices of thresholds for defining positive/negative interactions, as it only requires adjusting the corresponding thresholds and rerunning the code. For example, if we are interested in off-target predictions rather than the identification of primary targets, we can relax the activity threshold to a higher value of 10 $\mu$M to define positive DTIs. To illustrate this, we compute LCIdb_Orphan_10$\mu$M_threshold dataset in a manner similar to LDIdb_Orphan, but using a threshold of 10 $\mu$M rather than 100nM. The resulting LCIdb_Orphan_threshold dataset is much larger than LCIdb_Orphan and: it contains 1,037,934 positive DTI, 508,353 molecules, and 2,970 proteins. We focus on the orphan setting as this is the most challenging scenario.

Table S3 shows the performance of the Komet, ConPLex, MolTrans algorithms, as well as Random Forest on concatenated features, on the newly created LCIdb_Orphan_10$\mu$M _threshold. Komet still outperforms the other methods in the most challenging scenario (orphan), based on 5-fold cross-validation.

Table S3: Comparison of AUPR scores on LCIdb_Orphan_10$\mu$M_threshold, in 5-fold cross-validation

| Komet | ConPLex | MolTrans | RF with concatenated features |
|---|---|---|---|
| **0.862±0.012** | 0.838±0.012 | 0.555±0.004 | 0.606±0.006 |

### 5 Nyström approximation

In Komet, we encode molecules leveraging the Nyström approximation.[4,5] In the following, we present the mathematical details of Section 4.3.

Let us consider a set of landmark molecules $\{\hat{\mathfrak{m}}_1, \dots \hat{\mathfrak{m}}_{m_M}\}$, a new molecule $\mathfrak{m}$, and a kernel $k_M$ over molecules. The kernel matrix $K \in \mathbb{R}^{(m_M+1)\times(m_M+1)}$ over these $m_M + 1$ molecules can be written as $K = \begin{bmatrix} \hat{K}_M & \kappa^\top \\ \kappa & k_M(\mathfrak{m},\mathfrak{m}) \end{bmatrix}$ with $\hat{K}_M \in \mathbb{R}^{m_M\times m_M}$ being the kernel matrix over the landmark molecules and $\kappa = (k_M(\mathfrak{m}, \hat{\mathfrak{m}}_1), \dots, k_M(\mathfrak{m}, \hat{\mathfrak{m}}_{m_M})) \in \mathbb{R}^{m_M}$ the vector of kernel values between $\mathfrak{m}$ and the landmark molecules.

The Nyström's approximation consists in approximating $K$ as $K \approx C\hat{K}_M^{-1}C^\top = \begin{bmatrix} \hat{K}_M & \kappa^\top \\ \kappa & \kappa\hat{K}_M^{-1}\kappa^\top \end{bmatrix}$

with $C = \begin{bmatrix} \hat{K}_M \\ \kappa \end{bmatrix} \in \mathbb{R}^{(m_M+1)\times m_M}$.

Writing the Single Value Decomposition of $\hat{K}_M$ as $\hat{K}_M = U \operatorname{diag}(\sigma) U^\top$, the approximation of $K$ can be rewritten as $K \approx \Phi\Phi^\top$ with $\Phi = CU \operatorname{diag}(\sigma)^{-1/2} \approx CE$. When no dimensionality reduction is performed ($d_M = m_M$), $E = U \operatorname{diag}(\sigma)^{-1/2}$ and $\Phi = CE$.

The last line of matrix $\Phi$ is $\Phi_{m_M+1} = (\sum_{l=1}^{m_M} C_{m_M+1,l} E_{ls})_{s=1}^{m_M} = \psi_M(\mathfrak{m})$. Similarly, its $m_M$ first lines are $\psi_M(\hat{\mathfrak{m}}_1)$, ..., $\psi_M(\hat{\mathfrak{m}}_{m_M})$. Hence $k_M(\mathfrak{m}, \hat{\mathfrak{m}}_i) \approx \langle \psi_M(\mathfrak{m}), \psi_M(\hat{\mathfrak{m}}_i) \rangle$ for any molecule $\mathfrak{m}$ (including one of the landmark molecules), which justifies our proposition of $\psi_M$.

Furthermore, if we do not use dimensionality reduction, because the Nyström approximation is an equality on the upper-left block $\hat{K}_M$, $k_M(\hat{\mathfrak{m}}_i, \hat{\mathfrak{m}}_j) = \langle \psi_M(\hat{\mathfrak{m}}_i), \psi_M(\hat{\mathfrak{m}}_j) \rangle$ for any pair of landmark molecules.

### 6 Efficient computation

We explicit here the details for equality (a) of Equation 2 in Section 4.4.

$$(Zw)_k = \langle w, z_k \rangle_{\mathbb{R}^{d_Z}} \overset{(a)}{=} \langle m_{i_k}, W p_{j_k} \rangle_{\mathbb{R}^{d_M}} \overset{(b)}{=} \langle m_{i_k}, q_{j_k} \rangle_{\mathbb{R}^{d_M}}.$$

We use the matrix representation $W \in \mathbb{R}^{d_M \times d_P}$ instead of $w \in \mathbb{R}^{d_Z}$ in a way that $w$ is the flattened representation of $W$.

$$\begin{aligned} \forall k = 1..n_Z,\ (Zw)_k &= \langle w, z_k \rangle_{\mathbb{R}^{d_Z}} \\ &= \langle W, m_{i_k} p_{j_k}^\top \rangle_{\mathbb{R}^{d_M \times d_P}} \\ &= \operatorname{tr}\left( W (m_{i_k} p_{j_k}^\top)^\top \right) = \operatorname{tr}\left( W p_{j_k} m_{i_k}^\top \right) = \langle W p_{j_k}, m_{i_k} \rangle_{\mathbb{R}^{d_M}} \end{aligned}$$

### 7 Nearest Neighbor SVM model versus Komet

Komet uses a multi-task approach, that is to say, makes use of information about interactions that involve neither the query protein nor the query ligand. In order to showcase the benefit of such an approach, we focused on cases where the protein for which we want to predict ligands is orphan, that is to say, has no known ligands. In this setting, a single-task approach consists of training an SVM on a data set of molecules, which are labelled positive if they interact with the *nearest* protein (according to the LA kernel) to the query protein, and negative otherwise. We refer to this approach as single-task NN SVM, and compared it to Komet in several settings: on the whole LCIdb dataset, as well as on some specific cases: when working only within one family of proteins (either GPCRs or kinases), in cases where the nearest protein was close to the query protein (high similarity, defined by a LA kernel value greater than 0.75), or when it was far (low similarity, defined by a LA kernel value lower than 0.25).

For each protein in the considered data set considered in turn as the query protein, we performed the following experiment: a test set was built, comprising all known positive DTIs involving the query protein in LCIdb and their balanced negative DTIs. A corresponding training set is built: for Komet, it consists of all DTIs remaining in LCIdb after the removal of DTIs that are in the test set so that the query protein is orphan. The AUPR is calculated on the test set. For the NN SVM model, the training set consists of all positive and negative DTIs involving the nearest protein, according to the LAkernel. The AUPR is calculated on the test set, as for Komet.

Table S4 provides the mean and standard deviation of the AUPR for both approaches in all the aforementioned settings.

In all settings, Komet always shows better performance on average than the single-task NN SVM approach.

Table S4: Comparison of AUPR scores for Nearest Neighbor (NN) SVM and for Komet. Proteins with similar NN are those with which the nearest protein has a LA kernel greater than 0.75. Proteins with distant NN are those with which the nearest protein has a LA kernel lower than 0.25.

| | NN SVM | Komet |
|---|---|---|
| All proteins | 0.78±0.23 | **0.84±0.21** |
| Kinase superfamily | 0.80±0.22 | **0.92±0.12** |
| G-protein coupled receptor | 0.78±0.22 | **0.83±0.18** |
| Proteins with similar NN | 0.89±0.19 | **0.94±0.13** |
| Proteins with distant NN | 0.68±0.23 | **0.73±0.23** |
